# Exome Sequencing and Allele Dosage Analysis of Coast Redwood, a Hexaploid Conifer, Indicates Continuous Population Structure with a Population Break South of San Francisco Bay

**DOI:** 10.1101/2025.11.20.689601

**Authors:** Alexandra Sasha Nikolaeva, James Santangelo, Lydia Smith, Richard Dodd, Rasmus Nielsen

**Affiliations:** Department of Environmental Science, Policy and Management, University of California, Berkeley; Center for Computational Biology, University of California, Berkeley; Department of Integrative Biology, University of California, Berkeley; Museum of Vertebrate Zoology, University of California, Berkeley; Center for GeoGenetics, University of Copenhagen, Denmark

## Abstract

The coast redwood (*Sequoia sempervirens*) is a long-lived, hexaploid conifer of high ecological, cultural, and economic value whose range has been greatly reduced by historical logging [1]. Effective restoration and conservation depend on understanding patterns of genetic differentiation across the redwood range to delineate populations for management prioritization. Yet, past range-wide studies provided only a partial picture of population structure in coast redwood as they relied on a limited set of genetic markers [2, 3] or limited sampling, as sequencing was done on the same range-wide provenance collection [4, 5]. Here, we analyze 334,029 SNPs from a new range-wide set of 224 individuals using a dosage-based approach that accounts for polyploidy. Principal coordinates and clustering analyses reveal clear latitudinal structure, with a steeper break south of San Francisco Bay. Outlier SNP analysis identified new candidate loci involved in salinity tolerance, climate stress response, and nutrient uptake, suggesting potential local adaptation. These results point to the central role of geography in shaping genetic variation in coast redwood and give scientific basis for designing new conservation and management strategies aimed at preserving the species’ genetic diversity into the future.

## 2 Introduction

Coast redwood (*Sequoia sempervirens* (D. Don) Endl.) is a long-lived conifer that stretches along the coast of California, from southern Oregon to south of Monterey Bay. The species holds profound cultural significance for the Indigenous Peoples, including the Yurok, Karuk, Tolowa, and Coast Miwok Nations, who have inhabited these forests for millennia [6, 7]. Redwood was and still remains central to material culture of the Tribal Nations. In the past, split planks formed the walls of traditional houses, and fallen redwood logs were made into dugout canoes by hollowing it out to shape a canoe essential for salmon fishing on the Klamath and Smith rivers[6]. Indigenous Peoples also actively shaped redwood forest structures through cultural burning [8–10] and today this practice is returning to the redwood forests as one of the most important management tools.

Coast redwood is the tallest tree in the world [11], the state tree of California, and an unusual conifer in many respects. It is one of the few polyploid conifers [12, 13] and is considered an autohexaploid [14]. As is typical for polyploid species [15], it primarily reproduces asexually, though seed reproduction also plays a significant role in its population dynamics [16, 17]. Given its unique genetic and physiological characteristics and importance for conservation, researchers have naturally been drawn to study the species’ genetic diversity.

Understanding population structure is the typical first step towards understanding genetic diversity within any species. By comparing how allele frequencies in a species vary across populations, these studies can identify patterns of isolation and gene flow, which in turn provide insights into the overall population dynamics and adaptive potential of the species. The first attempt to incorporate full available genetic information for analysis of population structure in coast redwood was made in a 2011 paper by Douhovnikoff and Dodd [2]. They estimated genetic diversity among populations using microsatellite markers scaled by ploidy level. Their results indicated a weak divergence south of the Sonoma-Mendocino border and two or three population clusters as the most likely population structure. However, they conclude that a population break between northern and southern populations is located further north of the San Francisco Bay.

Later studies in the paternally inherited redwood chloroplast confirmed the San Francisco Bay as the boundary between southern and northern populations [3, 18]. Yet, in all previous studies of population structure, the number of markers was low (6 microsatellite markers in [2], 12 in [3]). In addition, previous studies relied on the same range-wide collection of redwood seed and cuttings made by Kuser [4], and then grown on the territory of the Russell Research Station in Lafayette, California. The most recent study on coast redwood [5], using exome sequencing and 64,358 SNPs on 92 individuals, was also conducted on the same collection. fastStructure analysis indicated *K* = 1 as the maximum marginal likelihood while *K* = 3 best explained the residual structure in the dataset; only five individuals were assigned to distinct population clusters, with the remaining 87 trees formed a single homogeneous group.

Studies on the phenotypic differences between the southern and northern redwoods suggest some variation in specific traits. Southern redwoods, for example, have a thicker bark than northern redwoods [19]. There were also significant differences in metabolic response to temperature among populations, with southern populations being more tolerant of higher temperatures [20]. However, in an experimental setting, only minor intraspecies differences in physiological responses to drought (shoot water potential, embolism, water use efficiency and photosynthesis) were found among populations [21].

The seed size also varies regionally, from the average of 4.7 mm in the north to 3.34 mm in length in the south (Claire Whicker, unpublished data) and similar observations of an increase in the seed size from south to north had been made earlier by the US Forest Service [22].

Summer fog is a key axis of environmental variation along the redwood range. Foliar water uptake and fog drip supply a substantial fraction of the tree’s summer water budget [23], and fog availability declines toward the southern range margin [23]. A high-resolution paleoclimate record from coastal northern California documents pronounced peaks in fog deposition at approximately 2,300 and 1,100 cal BP, inferred from diatom and silicoflagellate assemblages; these fog peaks coincide with increases in *Sequoia sempervirens* pollen [9, 24]. Repeated cycles of fog-driven expansion and contraction at the southern margin may have contributed to differentiation in osmotic and drought-stress loci, consistent with the high-*F_ST_* candidates we identify (discussed below).

Detecting population structure in polyploid plant populations presents both theoretical and computational challenges, as conventional methods, even though technically applicable to polyploids, require substantial modification and cautious interpretation [25]. Bayesian clustering approaches such as STRUCTURE, and more recent likelihood-based methods like ADMIXTURE, can be adapted for polyploids to estimate the number of populations (*K*) under assumptions of Hardy–Weinberg and linkage equilibrium [26–28], assuming accurate hexaploid genotype calling. However, accurate hexaploid genotype calling is challenging and requires very high sequencing depth. Principal components (PCA) and principal coordinate analyses (PCoA) is also widely used to explore population structure, but some implementations “diploidize” polyploid data, potentially distorting results [29–31]. By using allele dosages from the raw data, PCoA can be adapted to allele frequency matrices, while avoiding the biases introduced by forced diploidization. Another clustering alternative for polyploids is the LEA R package [32], which implements sparse non-negative matrix factorization (sNMF) directly on allele dosage data without requiring phased genotypes or explicit genotype calling. Another option is entropy [**Shastry2021MixedPloidy**], a Bayesian MCMC framework that estimates admixture proportions from genotype likelihoods and formally propagates genotype uncertainty; however, its computational demands scale steeply with the number of loci, making it prohibitive for genome-wide datasets with hundreds of thousands of SNPs. In this study, we use LEA sNMF as our primary clustering method, applied to allele dosage data to infer admixture proportions across the redwood range. We also use FreeBayes with hexaploid parameters to identify SNP positions from aligned reads (SNP calling), but calculate allele dosages directly from read counts to avoid the errors associated with genotype calling, resulting in (theoretically) unbiased estimates of quantities such as *F_ST_*. These dosage-based data were used to evaluate range-wide population structure and to perform an *F_ST_* -based genome scan to identify candidate loci potentially under selection.

## 3 Materials and Methods

### 3.1 Sample collection

We use the dataset previously described in [33]. These data were previously analyzed to infer patterns of chromosomal ploidy levels. Briefly, sampling was conducted in unmanaged old-growth and second-growth redwood forests, using the LEMMA forest structure dataset provided by the Save-the-Redwoods League. [34]. The history of the sampling site was confirmed with landowners, and while the exact age of trees was not determined, old-growth forests were estimated to be over 300 years old and second-growth trees were between 40-100 years old. Foliage samples, collected from the lowest branches or sprouts, were taken from trees at least 60 meters apart to avoid sampling clonal individuals. The location of each sample was recorded on Avenza maps (Avenza Maps™ v5.1.1). Samples were transported on ice to University of California, Berkeley campus, and stored at -80°C. A total of 305 tree samples were collected. The sequencing run included three pairs of technical replicates (trees sequenced in two independent libraries, samples 22 MRD/28 MRD, 13 286 1/3 286-1, and 16 416-1/35 416 1), which provided an empirical baseline for genotyping noise in our pipeline. Pairwise IBD distances within these replicate pairs were 0.085, 0.085, and 0.088, placing the empirical ceiling for clonal or replicate genetic identity at IBD *≈* 0.088. One sample of each technical replicate pair was removed. We additionally excluded all sample pairs whose pairwise IBD distance fell below this ceiling, and five close-relative pairs (pairwise distances 0.084–0.091; one sample per pair removed). Geographic coordinates were missing for some remaining samples, or replicate samples shared coordinates with trees from the same location, and these were also excluded. The minimum pairwise IBD distance among the 224 retained samples is 0.125 — approximately 40% above the highest technical replicate distance — confirming that no clonal samples or replicate remains in the dataset.

### 3.2 DNA extraction and sequencing

DNA was extracted from leaf tissue using a modified CTAB protocol, with additional steps including a chloroform pre-wash and ethanol wash. A magnetic bead clean-up was performed, and DNA purity and concentration were assessed. DNA was then diluted, sonicated, size-selected, and concentrated. Library preparation followed a modified Kapa Hyper Prep protocol, with adapters ligated and amplified. Libraries were combined into pools, hybridized using the Twist Target Enrichment Standard Hybridization protocol, and assessed for concentration. The final pool was sequenced across four lanes of Illumina NovaSeq X 10B, targeting 17.7Mb of conserved sequences selected from redwood genome annotations. We describe the full procedure of DNA extraction and sequencing in [33].

### 3.3 Read alignment and SNP calling

Raw reads were aligned to the reference genome published by The Redwood Genome Project (RGP) [35], using bwa mem [36] and SNPs were called on the BAM files. The pipeline is documented in a snakemake pipeline [https://github.com/James-S-Santangelo/prg]. We generated region files for Freebayes (v1.3.6) [37] using a custom Python script to divide the genome into regions of 1Mb for parallel processing.

We used Freebayes for SNP calling with the following parameters: –fasta-reference, –ploidy 6, –use-best-n-alleles 4, –haplotype-length -1, –max-complex-gap -1, –min-base-quality 20, –min-mapping-quality 30, –skip-coverage 50000, –targets, –bam-list, and –report-monomorphic. The SNP calling was conducted for each genomic region, and the resulting VCF files were concatenated using bcftools, followed by additional filtering to retain high-quality variants. SNPs were then filtered to include only bi-allelic sites using bcftools view. After filtering, we retained 334,029 bi-allelic SNPs. Mean sequencing depth per SNP was 17.70*×* with variance of 546.13. Notice that while this sequencing depth provides a good basis for estimating individual allele dosages, (i.e allele frequencies within an individual), it is not necessarily sufficient to provide accurate genotype calls in a hexaploid.

### 3.4 Genetic Distance Matrix

To estimate an Identity-By-Descent (IBD) distance matrix, we used the allelic counts from the sequencing reads. They were extracted from a compressed VCF file generated by Freebayes using the PyVCF Python library [38] to access the AD (Allelic Depth) field from the FORMAT column for each SNP. The allele dosage of the alternative allele in sample *i*, SNP *j* was then defined as

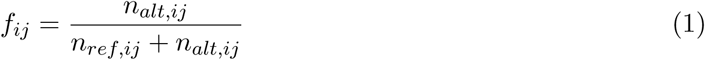

where *n_alt,ij_*and *n_ref,ij_*are the counts of the alternative and reference allele, respectively, in sample *i*, SNP *j*. If the AD field was missing or the total depth (*D*_ref_ +*D*_alt_) was zero, the dosage was recorded as missing. The resulting dosage values were stored in a matrix with samples as rows and SNPs as columns. To compute the genetic distance matrix, we used the Identity-By-Descent (IBD) distance, sometimes called the Identity-By-State (IBS) distance, but we do not distinguish between the two here. This distance estimates the probability that two alleles sampled from two different individuals are identical to each other. The advantage of using IBD distances in this setting is two-fold: (1) It is robust to low sequencing depth. The expected value of the statistic is a constant function of the sequencing depth so genomes with high and low coverage can be compared in an unbiased manner. (2) The distance has the same definition and statistical interpretation for any ploidy, i.e. for any ploidy it estimates the probability that two alleles sampled from the different individuals are different from each other. However, the biological interpretation may naturally differ depending on the setting. The IBD distance between two samples *i* and *j* is defined as

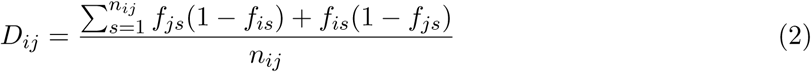

where *f_js_* and *f_is_* are the dosage values at the *s*-th SNP locus for samples *i* and *j*, respectively and *n_ij_* is the number of loci with non-missing dosage data for both samples. We exclude missing sites in order to correct for potential capture biases among samples. Notice that this distance estimates the same distance, in expectation, as the *p*-distance in phylogenetics, i.e. the uncorrected nucleotide divergence. When applied to two genetically identical individuals it estimates the heterozygosity, which naturally extends to polyploids through the concept of probability of IBD.

After initial filtering, we retained 334,029 bi-allelic SNPs, but to ensure complete coverage across all individuals, we further restricted the dataset to 139,067 SNPs with non-missing data in every sample. After that, we also refined the genetic distance matrix by adjusting for potential differences between PCR groups. The original distances were adjusted to account for the mean distances within and between PCR groups. The adjusted distance 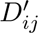 for samples *i* and *j*, belonging to PCR group *a* and *b*, respectively, was computed as

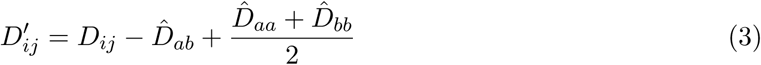

where *D_ij_* is the original distance between samples *i* and *j*; *D̂_ab_* is the mean distance between PCR groups *a* and *b*, and *D̂_aa_* and *D̂_bb_* are the mean distances within PCR groups *a* and *b*, respectively.

This step was necessary because we found a small bias in depth coverage among samples from different PCR groups, likely due to imperfect annotation sequences that overrepresented certain haplotypes. Each PCR group received variable number of PCR cycles, based on the initial DNA concentration of the samples (see [33] for full description of laboratory methods). The variation in depth coverage among PCR groups can affect the accuracy of genotype calls, potentially leading to biased allele frequency estimates and inaccurate representation of genetic diversity. To address this, we adjusted the genetic distances in hope to reflect true genetic differences rather than technical artifacts that might arise due to variable depth coverage. Full genetic distance matrix can be found in Supplementary Table S1. The origin and magnitude of the PCR-pool bias are illustrated in Appendix Figure A1, which shows PCoA with and without the correction, colour-coded by PCR pool group.

### 3.5 Principal Coordinates Analysis

We performed principal coordinates analysis (PCoA) on the genetic distance matrix to visualize variation among samples and explore potential associations with geographic latitude. The PCoA (classical multidimensional scaling) implementation was written in Python.

### 3.6 Calculating fixation index *F_ST_* from allele dosages

Based on the results of PCoA clustering, samples were divided into three populations using latitude boundaries: North (*≥* 40°N), Central (*≥* 38°N and *<* 40°N), and South (*<* 38°N). Allele frequencies were calculated separately for each population as well as for the full dataset. For each SNP, the allele frequency was estimated as the mean dosage across all individuals with non-missing data. This calculation was repeated within each population to obtain population-specific allele frequencies.

Then we calculated *F_ST_*for each SNP using the following approach:

First, the allele frequency *p_k_* was estimated for each population *k* as the mean dosage:

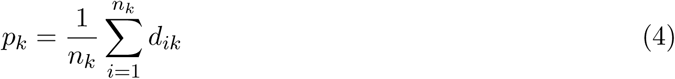

where: - *d_ik_*is the dosage (alternative allele frequency) of individual *i* in population *k*, - *n_k_*is the number of individuals with non-missing dosage in population *k*.

The total mean allele frequency *p̅* across all populations was computed as:

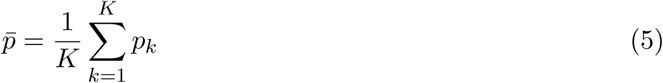

where *K* is the number of populations with valid data for the SNP.

We then calculated total heterozygosity, *H_T_*, and mean within-population heterozygosity, *H_S_*,

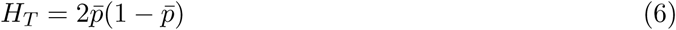

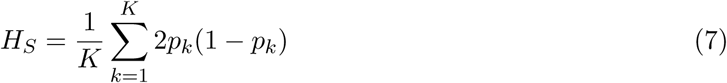

Finally, Wright’s fixation index *F_ST_* was calculated using the standard formula:

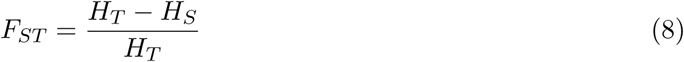

If *H_T_*= 0, *F_ST_*was not defined and recorded as missing. We note that this procedure for estimating *F_ST_*, by first estimating the allele frequency within populations using allelic counts pooling individuals without calling genotypes, has the advantage that it is robust to varying sequencing depth. We also note that we use a classical definition of *F_ST_*even though the organism we are working on is polyploid, because Wright’s *F_ST_*can be interpreted as analysis of probabilities of IBD that does not depend on the ploidy of the organism.

We then selected the top 1,000 SNPs with the highest *F_ST_* values. Flanking sequences (±300 bp) around each outlier SNP were extracted from the coast redwood reference genome using samtools faidx and aligned to the diploid reference genome of giant sequoia (*Sequoiadendron giganteum*; *SEGI*) using minimap2 [39] with short-read presets (-x sr --secondary=no) to establish chromosomal positions for visualization.

### 3.7 Population structure inference

To infer genome-wide admixture proportions, we used the sNMF function from the LEA R package [32] with ploidy set to 6. Input data consisted of allele dosage values for all 139,067 complete-case SNPs, rounded to the nearest integer (0–6 copies of the alternative allele). We ran sNMF for *K* = 2 to *K* = 5 with 10 independent repetitions per *K* and selected *K* using the minimum cross-entropy criterion. Admixture proportions (Q-matrices) at *K* = 2 and *K* = 3 were visualized as stacked bar plots ordered by latitude.

### 3.8 *F_ST_* -LFMM2 overlap and functional annotation

To prioritize candidate loci under potential local adaptation, we intersected the top-1,000 *F_ST_*outlier SNPs with SNPs significantly associated with at least one environmental variable (LFMM2, *q <* 0.05; see below). SNPs in both sets defined the *F_ST_ × LFMM2 overlap*. Flanking sequences (±300 bp) around each overlap SNP were queried against the UniProt/SwissProt database using diamond blastx [40] (e-value *≤* 10*^−^*^4^, minimum 35% amino-acid identity). For each overlap scaffold, we additionally selected the top-ranked non-overlap *F_ST_* outlier SNP and annotated it by the same BLASTx procedure. Hits from non-plant organisms with low identity were excluded, as were redundant hits already present in the overlap set.

### 3.9 Environmental association analysis (LFMM2)

Latent factor mixed model (LFMM2) analysis [41] was performed to identify SNPs significantly associated with environmental variables, using the lfmm2 function from the LEA package. We included 11 environmental predictors: 8 bioclimatic variables from CHELSA [42] (annual mean temperature BIO1, temperature seasonality BIO4, max temperature of warmest month BIO5, mean temperature of warmest quarter BIO10, annual precipitation BIO12, precipitation of driest month BIO14, precipitation seasonality BIO15, precipitation of warmest quarter BIO18), mean annual daytime fog frequency [43], distance to coast (km), and topsoil cation exchange capacity (0–5 cm) from SoilGrids [44]. Population structure was controlled by specifying *K* = 3 latent factors, consistent with the LEA sNMF results. Significant associations were identified at a Benjamini-Hochberg-corrected *q*-value threshold of 0.05 for each environmental variable independently. The variable with the lowest *q*-value was assigned as the primary environmental predictor for each significant SNP.

### 3.10 Isolation by distance and Mantel test

To test whether genetic differentiation is consistent with a pattern of isolation by distance (IBD), we performed a Mantel test [45] between the PCR-corrected IBD genetic distance matrix and a matrix of log-transformed great-circle geographic distances. Statistical significance was assessed using 9,999 permutations. A partial Mantel test was additionally performed conditioning on a binary matrix of population group membership (K=2 and K=3), to evaluate whether discrete population assignments explain any genetic variation beyond geographic distance. Both tests were performed in R using the vegan package [46].

### 3.11 Segmented regression

To identify discrete geographic transition zones along the latitudinal gradient, we fitted piecewise (segmented) linear regression models relating PCoA1 and PCoA2 to latitude [47]. For each axis, all candidate breakpoint latitudes were scanned and the value minimizing the residual sum of squares (RSS) was selected as the best-fit breakpoint. Significance was assessed by comparing the piecewise model to a simple linear model using an *F* -test. Ninety-five percent confidence intervals around the breakpoints were estimated by 999 bootstrap resamples of individuals.

## 4 Results

### 4.1 Principal Coordinates Analysis

The PCoA revealed clear genetic differentiation among the populations. The first two principal coordinates (PCoA1 and PCoA2) explained 2.74% and 1.14% of the total genetic variance, respectively (Figure 1). PCoA1 values correlated strongly with latitudinal and longitudinal gradients (Figure 3). The *R*^2^ value for Latitude vs. PCoA1 was 0.80, and the *R*^2^ value for Longitude vs. PCoA1 was 0.76. Samples from latitudes south of San Francisco Bay form a distinct cluster on the PCoA plot, indicating genetic differentiation from those collected further north. While the central and northern samples are more intermixed, there remains a discernible influence of latitude on genetic differentiation, as evidenced by the variation along the PCoA1 values. There were 5 samples that clustered closer to the central populations but were geographically from the southern populations. Two of these samples were located in San Francisco Peninsula, north of Santa Cruz, and three samples were found south of Monterey Bay (Figure 1).

**Figure 1:**
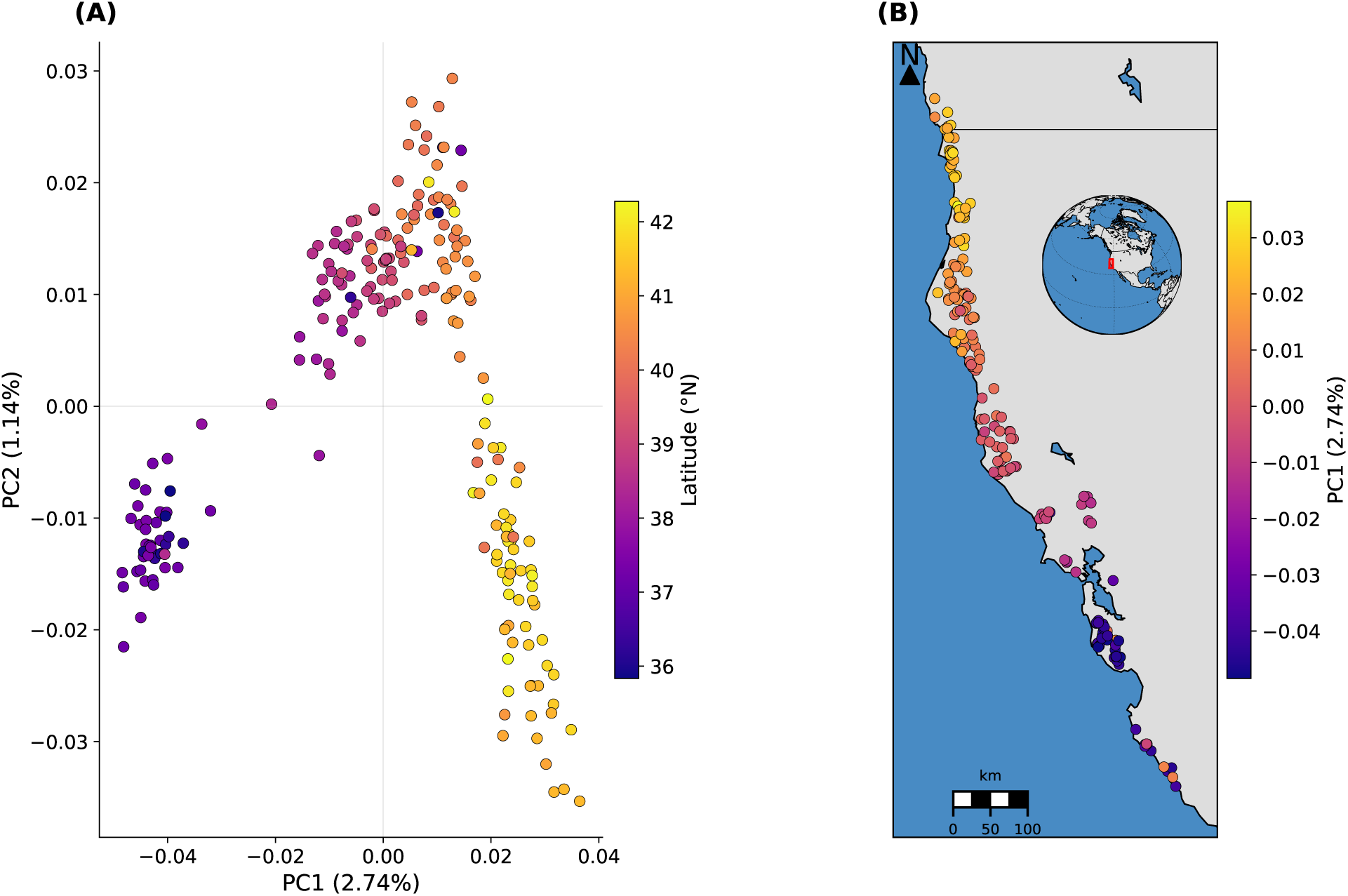
Genetic and geographic structure of coast redwood populations. **(A)** Principal Coordinates Analysis (PCoA) of the PCR-corrected IBD genetic distance matrix for 224 individuals, with points colored by latitude. PCoA1 (2.74% of variance) and PCoA2 (1.14% of variance) reveal a clear latitudinal gradient. **(B)** Geographic distribution of PCoA1 scores along the California and Oregon coast; warmer colors indicate higher PCoA1 values (southern individuals).

**Figure 2:**
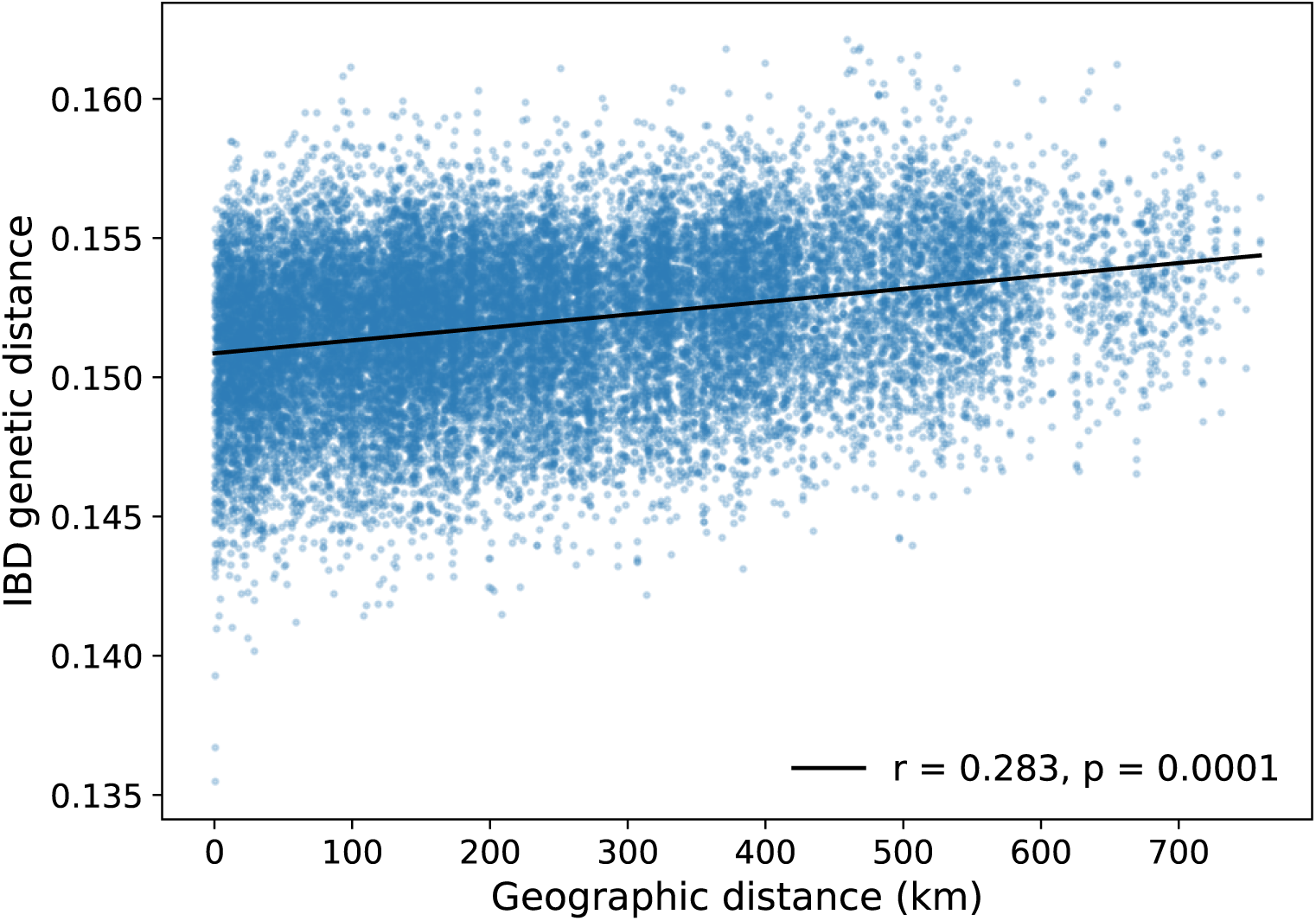
Isolation by distance in coast redwood. Scatter plot of pairwise IBD genetic distances against log-transformed geographic distances for all 224 sample pairs. *r* = 0.28, *p* = 0.0001 (9,999 permutations).

**Figure 3:**
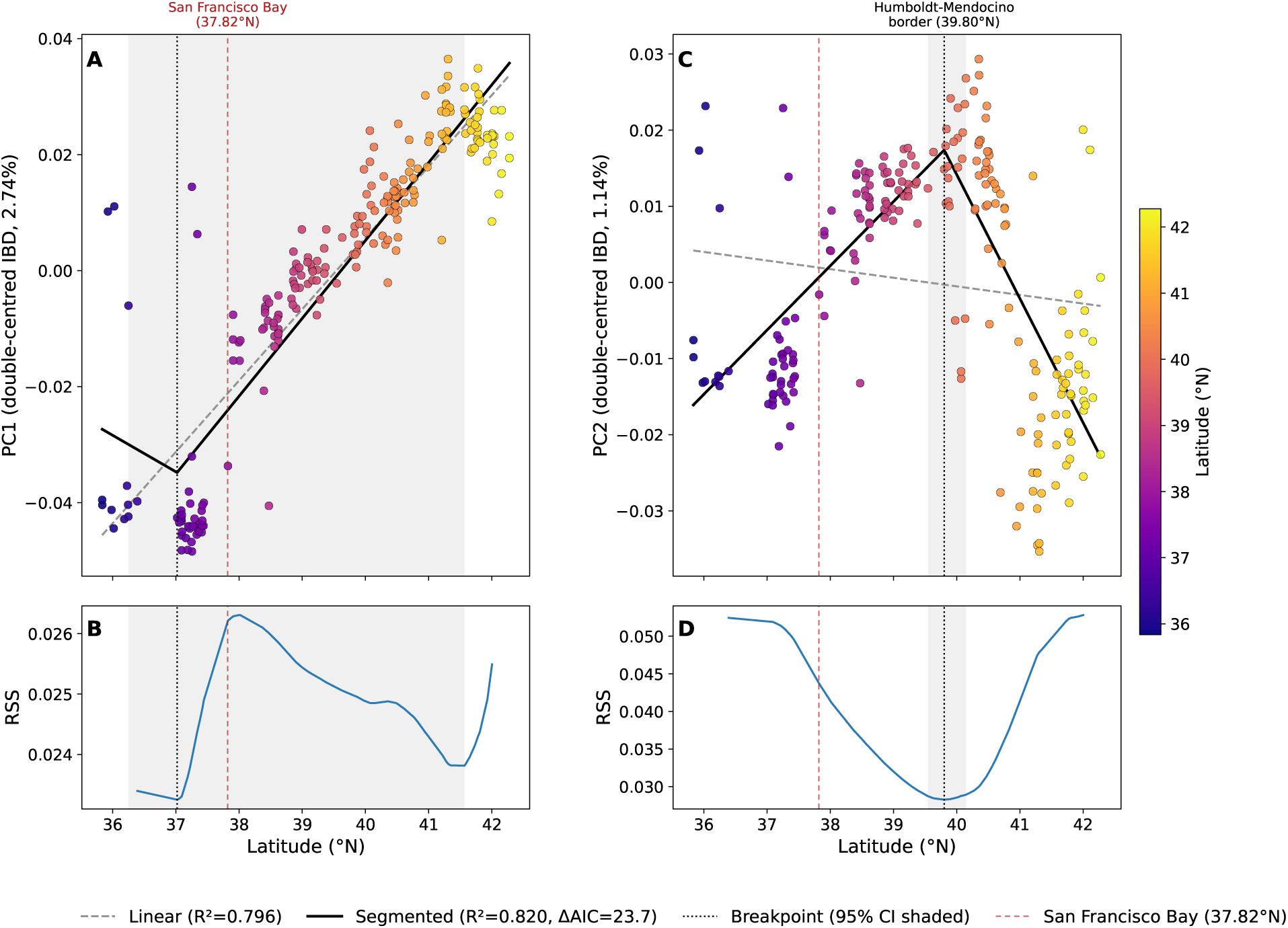
Segmented regression of PCoA axes on latitude for 224 coast redwood samples. **(A, C)** PCoA1 (2.74% of variance) and PCoA2 (1.14%) against latitude (°N), with individuals colored by region (North *≥*40°N, blue; Central 38–40°N, teal; South *<*38°N, orange). Solid black line = piecewise fit; dashed grey line = linear fit; vertical dotted line = best-fit breakpoint; shaded band = 95% bootstrap CI. Red dashed line in **(A)** marks San Francisco Bay (37.82°N); red dashed line in **(C)** marks the Humboldt–Mendocino county border (39.80°N). **(B, D)** Residual sum of squares (RSS) profile for all candidate breakpoint latitudes.

This geographic clustering remains consistent when PCoA is performed separately on old-growth and second-growth populations (Appendix Figure A2), although the population subdivision among the northern and central samples becomes less pronounced in the old-growth subset.

While we addressed PCR-pool batch effects in the calculation of the genetic distance matrix, minimal residual batch effects are reflected in PCoA2 values (Figure A1, Appendix A1).

### 4.2 Isolation by distance

A Mantel test between the PCR-corrected genetic distance matrix and log-transformed geographic distances revealed a significant positive correlation (Mantel *r* = 0.28, *p* = 0.0001, 9,999 permutations; Figure 2), confirming that isolation by distance is the dominant signal shaping genetic variation in coast redwood. Partial Mantel tests conditioning on population group membership (K=2 and K=3) yielded non-significant results (*p* = 0.75 and *p* = 0.84 respectively), indicating that once geographic distance is accounted for, discrete cluster assignment explains no additional genetic variance — consistent with a continuous clinal structure rather than discrete barriers.

### 4.3 Latitudinal transition zones

Piecewise linear regression of PCoA1 on latitude identified a significant breakpoint at 37.02°N (95% CI: 36.25–41.57°N; ΔAIC = 24.0, *F*_1,221_ = 29.5, *p <* 0.0001; Figure 3), corresponding to the San Francisco Bay region. This breakpoint marks where the latitudinal genetic gradient changes sharply, with southern populations showing higher PCoA1 values than northern and central populations. A second breakpoint in PCoA2 at 39.80°N (95% CI: 39.55–40.15°N; ΔAIC = 134.8, *F*_1,221_ = 189.7, *p <* 0.0001) corresponds to the boundary between central and northern populations near the Humboldt-Mendocino county border, where PCoA2 transitions from a positive to a negative slope. The narrow confidence interval for this northern breakpoint reflects the high sampling density in that region.

### 4.4 Admixture proportions

Non-negative matrix factorization with LEA sNMF (ploidy = 6) recovered minimum cross-entropy at *K* = 3, consistent with the three geographic clusters identified by PCoA and the two latitudinal breakpoints from segmented regression. At *K* = 2, nearly all southern samples form a distinct ancestry component, when as northern and central individuals are mostly assigned to a shared ancestry (Figure 4). At *K* = 3, a distinct northern cluster emerges above *≈*41°N, broadly consistent with the PCoA2 breakpoint near the Humboldt–Mendocino county border (39.80°N). Central samples show the highest degree of admixture between the northern and southern populations.

**Figure 4:**
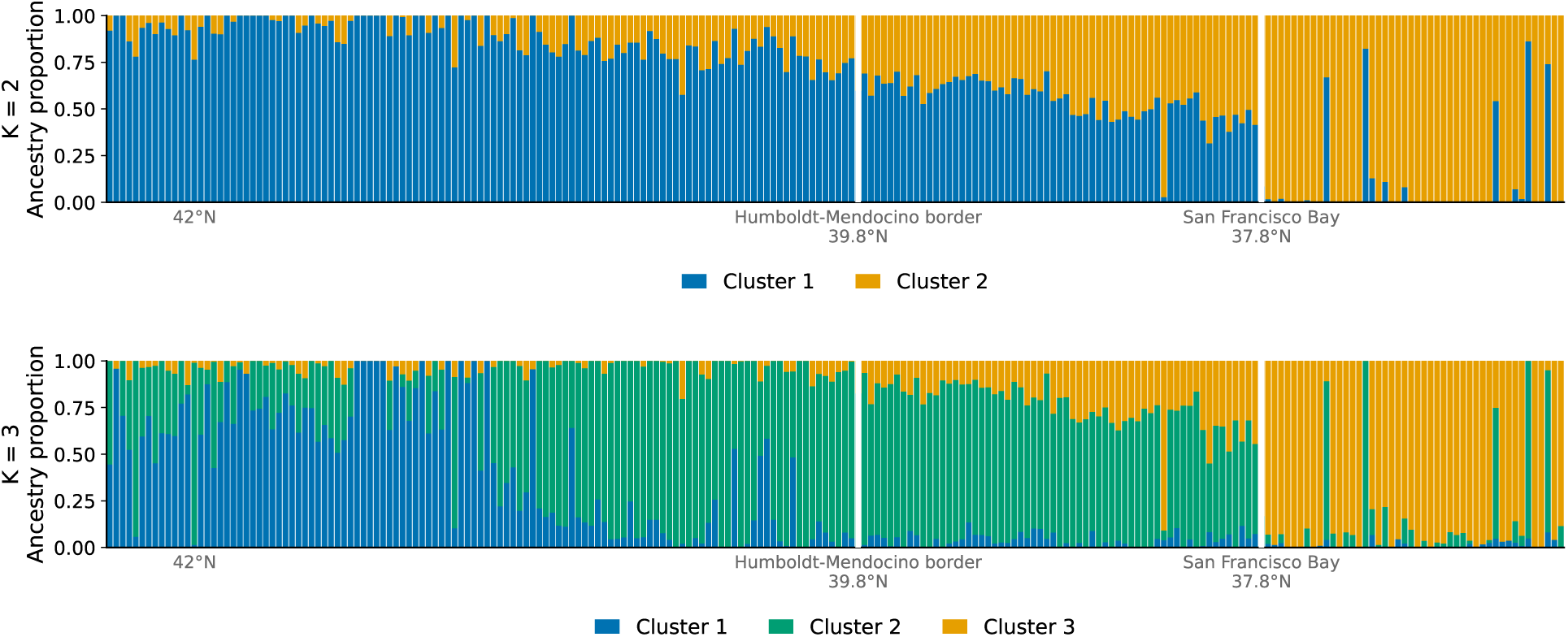
LEA sNMF admixture proportions for 224 coast redwood individuals at *K* = 2 (top) and *K* = 3 (bottom), ordered by latitude (north to south, left to right). White vertical lines mark San Francisco Bay at the Golden Gate Bridge (37.82°N) and the Humboldt–Mendocino county border (39.80°N), corresponding to the PC1 and PC2 segmented regression breakpoints respectively.

### 4.5 Overlap between ***F_ST_*** outliers and LFMM2 environmental association SNPs

To identify SNPs with the strongest differentiation among populations and potential signals of local adaptation, we calculated per-SNP *F_ST_* values across the three populations and selected the top 1,000 outlier SNPs. We then intersected these *F_ST_* outliers with SNPs showing significant environmental associations (LFMM2, *q <* 0.05, *K* = 3 latent factors), yielding 58 overlap SNPs distributed across multiple giant sequoia chromosomes (Figure 5).

**Figure 5:**
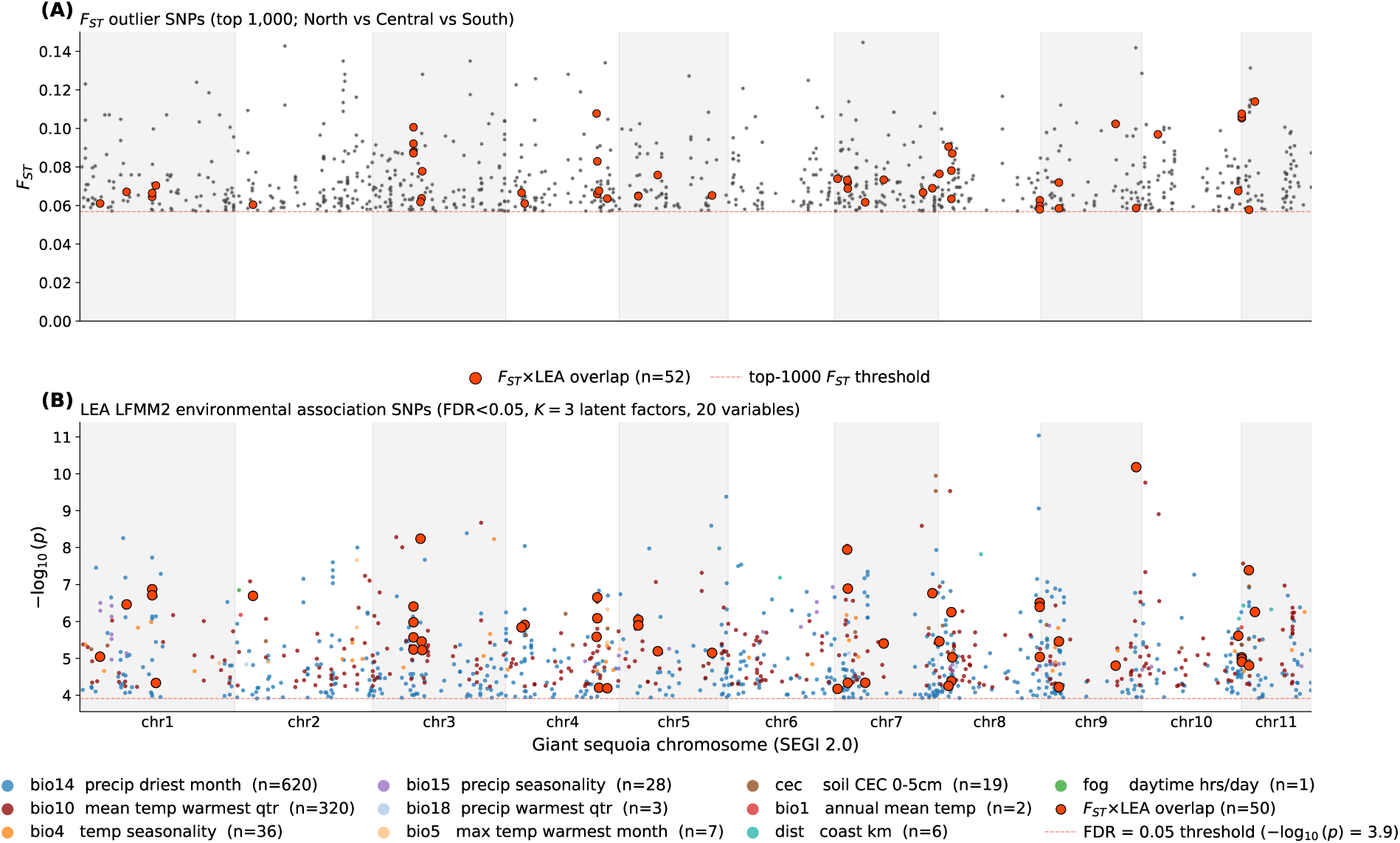
Combined Manhattan plot showing *F_ST_* outliers and LFMM2 environmental associations in coast redwood, mapped to the giant sequoia (*SEGI 2.0*) reference genome. **(A)**: per-SNP *F_ST_* values for the top-1,000 outliers comparing North (N), Central (C), and South (S) populations (grey points); orange points are SNPs also significant in LFMM2 (*q <* 0.05, *K* = 3). **(B)**: LFMM2 *−* log_10_(*p*) values for all significantly associated SNPs, colored by the environmental variable with the strongest association (PDM = precipitation of driest month; MAT = mean annual temperature; CEC = cation exchange capacity (used as a proxy for groundwater salinity)). Chromosomes 1–11 of the *SEGI* genome are shown along the x-axis.

Beyond these 58 exact-match overlap SNPs, we identified an additional 359 *F_ST_*outlier SNPs and 362 LFMM2-significant SNPs that are co-localized within 1-Mb windows across 211 genomic regions on all 11 giant sequoia chromosomes, where different SNP positions within the same region independently show both high *F_ST_*and significant environmental association (visible as coincident but non-orange dots in both panels of Figure 5; Supplementary Table S4).

To investigate the potential functional relevance of the most highly differentiated SNPs, we annotated all 58 overlap SNPs by BLASTx against UniProt/SwissProt (e-value *≤* 10*^−^*^4^, identity *≥* 35%); 52 returned a significant hit and are listed in Supplementary Table S3, while the remaining 6 had no match in the SwissProt database. Table 1 summarizes the top 10 candidate loci by *F_ST_*. The full list of *F_ST_* values can be found in Supplementary Table S2.

**Table 1:**
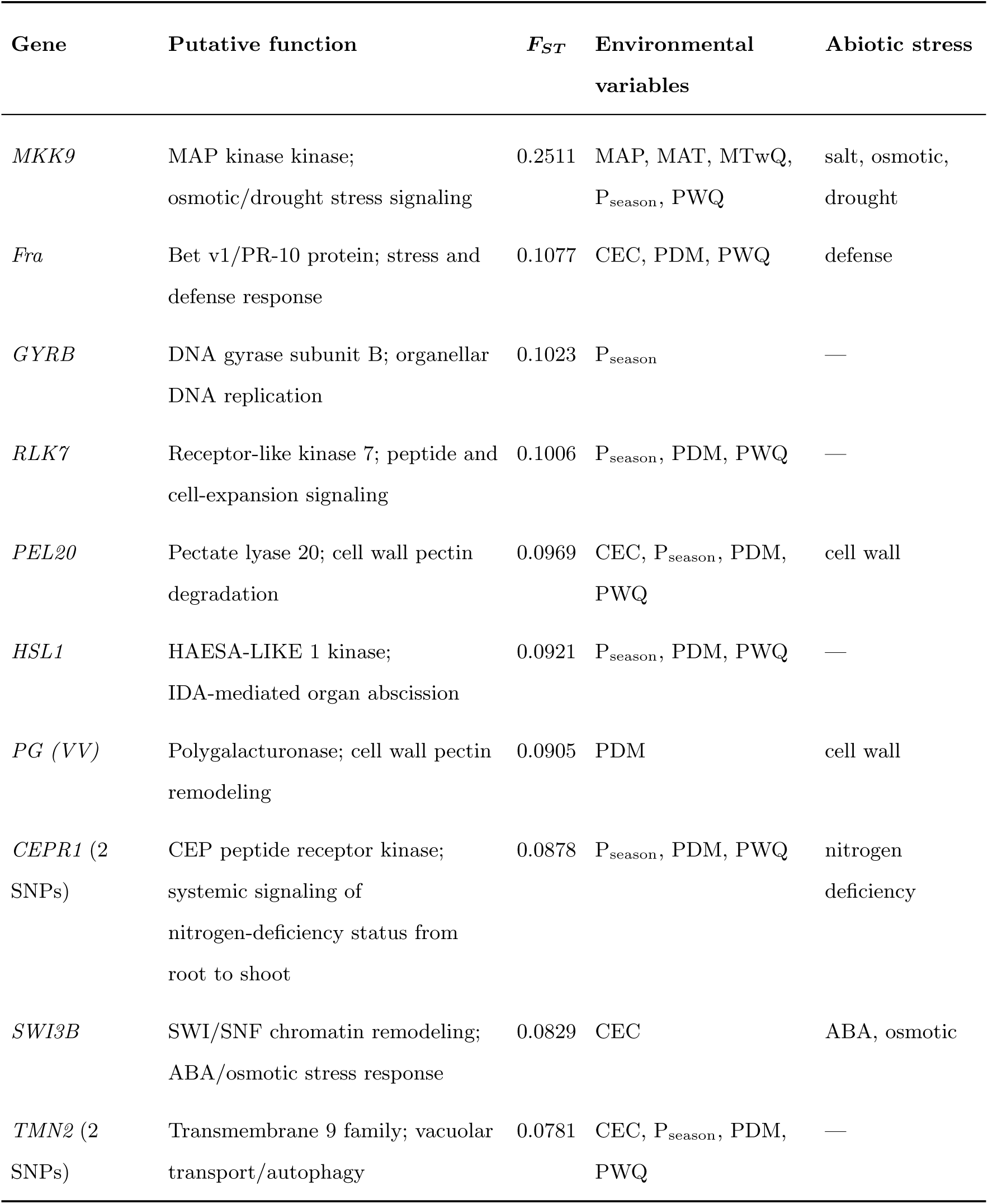
Top 10 *F_ST_* outlier loci at the intersection with LFMM2 environmental associations (top-1,000 *F_ST_* outliers *∩* LFMM2 *q <* 0.05, K = 3 latent factors), annotated by BLASTx against UniProt/SwissProt. Env. variables: MAT = mean annual temperature, MTwQ = mean temp. of warmest quarter, MAP = mean annual precipitation, PDM = precip. of driest month, P_season_ = precipitation seasonality, PWQ = precip. of warmest quarter, CEC = cation exchange capacity (0–5 cm). Full list in Supplementary Table S3.

| Gene | Putative function | $F_{ST}$ | Environmental variables | Abiotic stress |
| --- | --- | --- | --- | --- |
| *MKK9* | MAP kinase kinase; osmotic/drought stress signaling | 0.2511 | MAP, MAT, MTwQ, $P_{\text{season}}$, PWQ | salt, osmotic, drought |
| *Fra* | Bet v1/PR-10 protein; stress and defense response | 0.1077 | CEC, PDM, PWQ | defense |
| *GYRB* | DNA gyrase subunit B; organellar DNA replication | 0.1023 | $P_{\text{season}}$ | — |
| *RLK7* | Receptor-like kinase 7; peptide and cell-expansion signaling | 0.1006 | $P_{\text{season}}$, PDM, PWQ | — |
| *PEL20* | Pectate lyase 20; cell wall pectin degradation | 0.0969 | CEC, $P_{\text{season}}$, PDM, PWQ | cell wall |
| *HSL1* | HAESA-LIKE 1 kinase; IDA-mediated organ abscission | 0.0921 | $P_{\text{season}}$, PDM, PWQ | — |
| *PG (VV)* | Polygalacturonase; cell wall pectin remodeling | 0.0905 | PDM | cell wall |
| *CEPR1* (2 SNPs) | CEP peptide receptor kinase; systemic signaling of nitrogen-deficiency status from root to shoot | 0.0878 | $P_{\text{season}}$, PDM, PWQ | nitrogen deficiency |
| *SWI3B* | SWI/SNF chromatin remodeling; ABA/osmotic stress response | 0.0829 | CEC | ABA, osmotic |
| *TMN2* (2 SNPs) | Transmembrane 9 family; vacuolar transport/autophagy | 0.0781 | CEC, $P_{\text{season}}$, PDM, PWQ | — |

## 5 Discussion

In this paper we revisit questions of population structure in coast redwood, a hexaploid and clonal conifer, and investigate alleles that might point to local adaptation among the three redwood populations. Our results suggest that the southern redwood population is genetically distinct, which warrants special considerations for its conservation and management.

Separating neutral processes such as genetic drift or isolation-by-distance from the effects that arose due to local adaptation (or isolation-by-environment) is difficult even in diploid populations [48], but it is a necessary step in the search for signals of selection in natural populations. In this paper, we used PCoA and LEA sNMF to identify genetic structure, which informed our geographic delineation of redwood populations. We then treated these populations separately to detect SNPs that may have been subject to local adaptation within individual populations.

### 5.1 Geography plays a role in redwood population differentiation

Our results provide strong evidence that geography plays a significant role in the genetic differentiation of coast redwood populations. The Principal Coordinates Analysis (PCoA) performed on the genetic distance matrix showed that the first principal coordinate (PCoA1) explains a significant portion of the genetic variation (2.74%), with clear correlations to latitude.

Samples from southern populations, particularly those located south of San Francisco Bay, formed a cluster, suggesting some geographic influence on genetic structure. It is not exactly clear if this clustering is solely driven by geographic distance or there are other processes that might be contributing to a stronger differentiation among the southern population and the rest of the range. However, the existence of the southern cluster in our data supports earlier findings based on chloroplast data [3, 18], which identified the San Francisco Bay as a significant geographic barrier. Beyond the southern cluster, PCoA and LEA sNMF both indicate the existence of northern and central population groupings. A northern cluster is visible in the PCoA above approximately 41°N — just above Arcata Bay (specifically at Patrick’s Point, around 41.10°N) — though it is less pronounced in the old-growth subset (Appendix Figure A2), consistent with lower old-growth sampling density in the north. At *K* = 3, LEA sNMF identifies a distinct ancestral component for these northern samples, broadly concordant with the PCoA2 breakpoint at 39.80°N. The central cluster spans the region between the Humboldt–Mendocino county border and the Mendocino–Sonoma county border, which was previously identified as a boundary to gene flow between southern and central redwood populations [2]. However, a gap in our sampling in that area may have influenced the PCoA results, which are sensitive to uneven coverage [49], and the apparent discontinuity near the Sonoma–Mendocino border may partly reflect limited sampling rather than a true barrier.

Interestingly, five samples from second-growth populations in the southern part of the range cluster with northern samples from northern Mendocino (Figure 1). This could indicate possible gene flow from northern populations to southern latitudes. However, it seems more likely that these samples were planted in the southern part of the range. Contamination or accidental sample mislabeling cannot be ruled out either.

### Calculation of ***F_ST_*** from Mean Allele Dosage

In our study, we calculated *F_ST_* values using a dosage matrix, which provides fine-scale resolution of allele frequency differences. We use allele dosage rather than genotype calls because of the high genotyping uncertainty. For each SNP, we estimated population allele frequencies as the mean dosage across individuals. This circumvents the need for genotype calling and provides an unbiased estimator. We then used these population-level means to calculate expected heterozygosity within populations (*H_S_*) and across the total sample (*H_T_*) using standard formulas, which depend solely on allele frequencies: *H* = 2*p*(1 *− p*). This approach can be used because of the fact that expected heterozygosity is a function of population-level allele frequency and allows us to estimate *F_ST_* in a way that preserves signals of population structure independent of ploidy. One might argue that *H_T_* and *H_S_* calculated using the formula 2*p*(1 *− p*) relates to diploid Hardy-Weinberg expectations. However, these statistic estimate probabilities of IBD within the total population and the sub-populations, respectively, and are equally interpretable for polyploid and for diploid organisms.

### 5.2 Signals of local adaptation

Our results identify several candidate loci that may be under selection among redwood populations. The highest observed *F_ST_* value (0.251) was associated with a SNP located in a genomic region putatively involved in osmotic and drought stress signaling. This SNP falls within a sequence homologous to a predicted gene *MKK9* (Mitogen-Activated Protein Kinase Kinase 9)-like in *Cryptomeria japonica*, a key regulator in abiotic stress response pathways. In *Arabidopsis thaliana*, *MKK9* activates downstream MAP kinases such as MPK3 and MPK6 and plays a role in ethylene signaling and leaf senescence. Notably, *mkk9* loss-of-function mutants show increased tolerance to salt and osmotic stress during germination, suggesting that *MKK9* acts as a negative regulator of stress-responsive gene expression [50]. Previous studies on redwood adaptation have also highlighted differentiation in MAPK genes [5]; however, our results pinpoint a putative SNP within MKK9, a specific upstream kinase in the MAPK cascade that activates MPK3 and MPK6.

Additional *F_ST_* outliers co-occurring with LFMM2-significant loci are annotated to genes involved in cell wall remodeling (pectate lyase, polygalacturonase), chromatin remodeling (SWI3B), receptor kinases (RLK7, CEPR1), iron redistribution (LPR), and organellar functions (GYRB, PPR), consistent with broad environmental differentiation across precipitation, temperature, and soil gradients (Table 1; Supplementary Table S3).

Previous studies have suggested that, despite its distribution near the Pacific Ocean, coast redwood does not tolerate high levels of soil salinity or prolonged exposure to salt-laden winds. Greenhouse experiments by Nackley and colleagues [51] investigated the effects of different salts on the growth of coast redwood cultivar ‘Aptos Blue’. Their results showed that growth was significantly reduced when salinity in irrigation water exceeded 3 dS m*^−^*^1^. Diameter growth was especially sensitive, with 72–82% of the variation in trunk diameter explained by increasing salinity, while stem height showed a weaker correlation. Another study by Wu and Guo [52] confirmed that coast redwood is sensitive to foliar salt exposure. They observed visible stress symptoms such as needle burn, chlorosis, and biomass reduction when trees were sprayed with salt and boron solutions. Their results suggest that even moderate levels of salt spray can negatively affect coast redwood. However, both of these studies were conducted on Aptos Blue and Los Altos redwood cultivars, which originated from the southern redwood population. Currently, there are no studies on the range-wide salt tolerance in coast redwood and it is not clear whether northern populations demonstrate the same response to salt exposure as southern populations.

It is also likely that, rather than mediating response to salinity, MKK9-like genes are involved in broader osmotic stress signaling pathways. In *Arabidopsis*, MKK9 acts via the MKK9–MPK3/MPK6 module to promote leaf senescence, with loss-of-function mutants showing delayed senescence and overexpression causing premature aging [53]. This is notable because water-stressed redwoods have been observed to experience needle yellowing and branch die-back [54]. This kinase cascade also intersects with ethylene and ABA signaling pathways, influencing EIN3-dependent transcription and the coordination of stress responses [53, 55]. MAPK cascades (e.g., MKK1/MKK2–MPK6) have been shown to regulate stomatal closure via ABA signaling and control ROS and ion channel activity under osmotic or salt stress [56]. Genetic variation in these signaling components could allow redwood populations to fine-tune physiological responses to drought, fog, or other water-related stressors across their range.

Another notable outlier is a SNP located on Scaffold 387868 (HRSCAF=412686 139448964; Supplementary Table S3), which appears to be a distant homolog of a predicted multicopper oxidase gene, potentially related to the *LPR* (Low Phosphate Root) gene family. In *Arabidopsis thaliana*, members of the *LPR* gene family regulate multicopper oxidases involved in root development and phosphate homeostasis under nutrient stress [57]. LPR proteins are also implicated in iron redistribution, suggesting that the observed SNP may lie in a regulatory region affecting nutrient-responsive developmental processes or oxidative stress adaptation.

Foliar nutrient data from [58], (Table 4), indicate that coast redwoods growing on young, fertile alluvial or alluvial-derived soils accumulate high levels of nitrogen, phosphorus, calcium, magnesium, potassium, sodium and iron. By contrast, trees growing on highly weathered ultisols, which are high in iron oxide that is not easily available to plants, show deficiencies in nitrogen, calcium, magnesium and substantially reduced foliar phosphorus [58].

In the coast redwood region, ultisols are predominantly found on ridge tops and upland slopes [58]. At the geographic extremes of the redwood range, particularly toward the southern limit, trees show imbalances in foliar nutrient concentrations, including low iron levels and, in some cases, manganese deficiencies [58]. Geochemical data from the USGS soil survey indicate a north-to-south decline in total iron concentrations in surface soils across the coast redwood range, with some of the lowest iron levels observed in the San Francisco Bay Area and adjacent southern sites [59].

Interestingly, at greater tree heights, foliar iron concentrations are low, and a morphological transition is observed when redwood foliage shifts from the typical flat “peripheral” leaves to scale-like “axial” leaves resembling those of *Sequoiadendron giganteum* [58]. The proportion of axial foliage is higher in southern redwood populations compared to northern ones [60]. This shift has been interpreted as an adaptation to reduced fog and moisture availability in southern regions, allowing trees to regulate water uptake under hydraulic constraints. We suggest that, in addition to moisture stress, this shift may also reflect geographic differences in iron availability, which can affect photosynthesis. In *Arabidopsis*, the enzymes *LPR1* and *LPR2* help convert Fe(II) to Fe(III), supporting iron transport to leaves [61].

Due to the complexity of the hexaploid genome and the coexistence of both disomic and multisomic inheritance in coast redwood [62], patterns of linkage disequilibrium (LD) are expected to be varying across the genome. LD may decay more rapidly under multisomic inheritance, because frequent recombination among all homologous chromosomes accelerates the breakdown of linkage. In contrast, regions with preferential or disomic pairing may retain LD over longer distances. Therefore, the association between candidate outlier SNPs and functional loci under selection must be interpreted with caution.

### 5.3 Conservation and management implications for coast redwood

One of the key goals in coast redwood conservation is to restore second-growth redwood stands to an old-growth structure [63]. However, this goal may be difficult to achieve, as old-growth forests developed under different climatic conditions than those present today and in the future. The more urgent priority, though, is to ensure the long-term stability of the population and to preserve the existing genetic diversity, while maintaining it’s role as a valuable timber resource for the community. Our results suggest that the geographically isolated southern population south of the San Francisco Bay warrants prioritization in conservation planning. Historical logging removed approximately 96% of old-growth coast redwood [64], and the southern range is both more fragmented and more exposed to projected reductions in fog frequency and summer precipitation. Preserving genetic diversity in this population should be an explicit management objective.

There are also different management strategies for adapting forest trees to the effects of climate change, one of them being assisted migration [65]. This strategy involves relocating individuals with “adapted” genotypes from areas of drier or hotter climate into habitats that over time will become more suitable to that genotype.

The continuous structure we document here, with a more pronounced break around the San Francisco Bay suggests that the gene flow between the southern population and the rest of the range might have been limited, even though southern ancestry is present to some degree in the rest of the population. It would be necessary, therefore, to evaluate how individuals and clones from different parts of the range perform in a reciprocal transplant experiment. Such an experiment would establish whether the genetic breaks might also correspond to fitness differences under common environments and would clarify whether this population structure reflects local adaptation as well.

Our LFMM2 results suggest differences in how trees from different redwood populations might respond to changes in temperature, precipitation, and groundwater salinity. Therefore, we do not want to follow the assumption that species might be able to simply shift northward as the climate warms: broader evidence shows that forest species ranges can shift in multidirectional and sometimes unexpected directions, driven by the interplay of multiple environmental factors, including atmospheric nitrogen deposition [66, 67]. The genotype-environment associations we identify with LFMM2 and with with *F_ST_* scans provide a baseline for genomic offset modeling, which can quantify population-level vulnerability to projected climate change. However, such predictions are highly method-dependent and require validation against independent mortality or growth data before informing seed transfer decisions [68].

### 5.4 Study limitations

One of the major challenges in this study was the incomplete and imperfect annotation of the draft reference genome, which may have introduced bias in exome capture. We used annotation sequences from the published draft reference genome for coast redwood [35], but it was unclear whether the exome probes were originating from multiple haplotypes or a single haplotype. Such uncertainty can affect capture efficiency, as imperfect annotations may cause certain haplotypes or repetitive regions to be over- or underrepresented. Even minor differences in the capture protocol could exacerbate these effects.

Bias from non-uniform exome capture amplification during library preparation differs from that caused by PCR duplicates, which involve multiple copies of the same template molecule [69]. In this case, variation in the copy number of target probes themselves (i.e., identical sequences present on different chromosomes or duplicate sequences present on the same chromosome) can lead to sequences not being flagged as duplicates during downstream analysis, since reads from different haplotypes or different regions may carry different adapters before capture. If the probes were not deduplicated prior to synthesis, only some haplotype sequences may be captured, further distorting representation.

This uneven amplification can cause certain regions to be underrepresented or overrepresented, particularly if minor differences occur during library preparation. We detected such biases in our dataset and accounted for them when calculating the genetic distance matrix. However, bioinformatic correction is not a complete solution: extra copies of target probes can still lead to overrepresentation of specific alleles from individual chromosomes or haplotypes, producing inaccurate dosage estimates. This is especially problematic in polyploid species, where such biases can misrepresent allelic diversity and lead to incorrect conclusions about genetic variation, particularly in smaller datasets.

Future work should incorporate more careful probe design, deduplication of probes, and removal of those likely to amplify repetitive elements such as transposable elements or plastid-derived sequences. These steps would help reduce bias and lower sequencing costs.

A second limitation is the use of the diploid giant sequoia (*SEGI* 2.0) genome as a heterologous reference for mapping *F_ST_* and LFMM2 outlier SNPs to chromosomal positions. While minimap2 alignment of ±300 bp flanking sequences provides reasonable synteny-based positions, cross-species alignment introduces positional uncertainty, particularly in repeat-rich or rapidly evolving regions. Functional annotations were inferred from BLASTx similarity to plant proteins in UniProt/SwissProt rather than from direct annotation of the coast redwood genome, and gene identities should be treated as putative homologs.

A third limitation concerns the application of LFMM2, which was developed and validated primarily for diploid organisms. Although we supplied allele dosage values as input and the latent factor framework is not strictly ploidy-dependent, the calibration of *p*-values and *q*-values under hexaploidy has not been formally established. Our environmental association results should therefore be interpreted as exploratory signals rather than confirmed loci under selection.

## 6 Conclusions

This study revisited the question of neutral population structure in coast redwood (*Sequoia sempervirens*) using exome sequencing data from a new range-wide collection of 224 individuals. Our results suggest a clinal genetic structure shaped by isolation by distance (Mantel *r* = 0.28, *p* = 0.0001), with a pronounced steepening of the latitudinal gradient south of San Francisco Bay. PCoA, segmented regression, and LEA sNMF consistently identify this southern break as the most significant genetic discontinuity in the range, with a secondary transition near the Humboldt–Mendocino county border (39.80°N) marking the northern population boundary.

Intersection of *F_ST_* outliers with LFMM2 environmental associations identified 58 candidate loci, of which 52 were annotated to genes involved in osmotic stress signaling, nutrient uptake, lignin biosynthesis, and cell wall remodeling — consistent with adaptive differentiation along precipitation, temperature, and soil gradients across the range.

Our results provide scientific basis for prioritizing the geographically isolated southern population in conservation planning. We strongly advocate for range-wide reciprocal transplant experiments to evaluate performance across the identified populations, and for provenance selection strategies that account for limited gene flow across the San Francisco Bay.

This study also demonstrates that dosage-based polyploid methods can successfully recover population structure and adaptation signals in a hexaploid conifer. We encourage similar approaches in other polyploid forest trees where conventional diploid methods may not fully capture existing genetic diversity.

## 8 Data Availability

The raw sequencing data generated in this study have been previously deposited in the NCBI Sequence Read Archive (BioProject: PRJNA1163354). The data are publicly accessible and can be freely downloaded for academic and research purposes. Source code is available at https://github.com/SashaNikolaeva/Coast Redwood Population Structure

## Acknowledgements

The authors thank undergraduate researchers Liam Galleher, Claire Whicker, Simone Stevens, Nic Dutch, and Jenifer Camarena for their assistance in sample collection and DNA extraction. We also thank Michelle Davila for help in library preparation. We are also grateful to participating land managers who provided access to sampling locations, including those at The Napa Valley Reserve (Paul Asmuth), California State Parks, The California Department of Forestry and Fire Protection(CalFire) and especially Lynn Webb, Green Diamond Resource Company (Carlos Gantz and Scott Whittington), Mendocino Redwood Company, Humboldt Redwood Company, and The Lyme Timber Company.

## Appendix A

**Figure A1:**
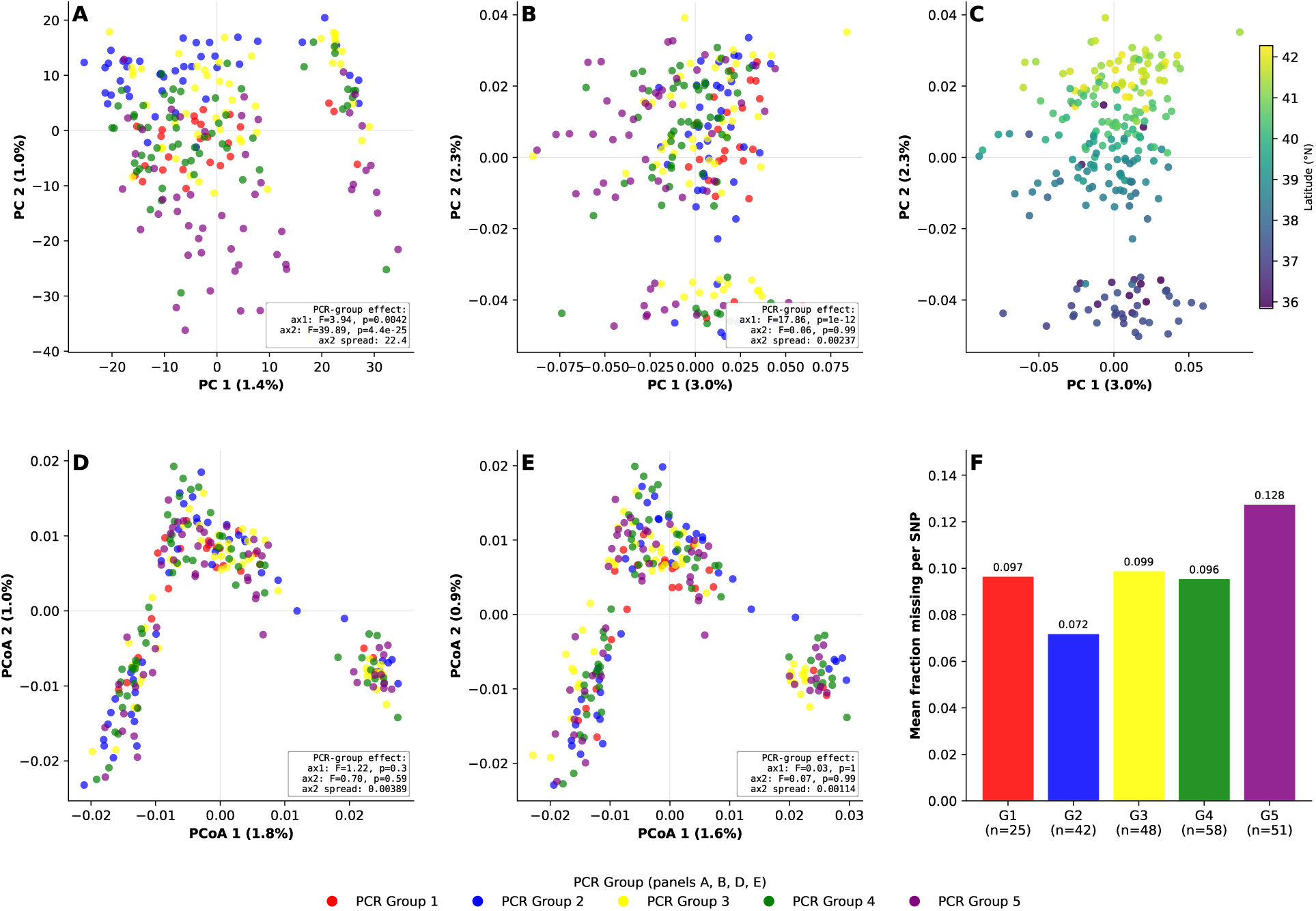
Assessment of PCR-pool batch effects on population structure inference. **(A)** PCA on the uncorrected allele dosage matrix (PC1/PC2), colored by PCR pool group; the pool-driven separation establishes the bias. **(L)** sklearn PCA on the dosage-corrected IBD distance matrix, colored by PCR pool group; residual pool separation is markedly reduced. **(H)** Same dosage-corrected IBD matrix as in (L), colored by latitude; the north-to-south gradient is preserved after correction. **(C)** PCoA on the uncorrected IBD distance matrix, colored by PCR pool group. **(D)** PCoA on the dosage-corrected IBD distance matrix, colored by PCR pool group; the pool effect is minimal. **(E)** Mean per-SNP missing-data fraction by PCR pool group, showing the origin of the bias (groups with higher missing-data rates received fewer PCR cycles).

**Figure A2:**
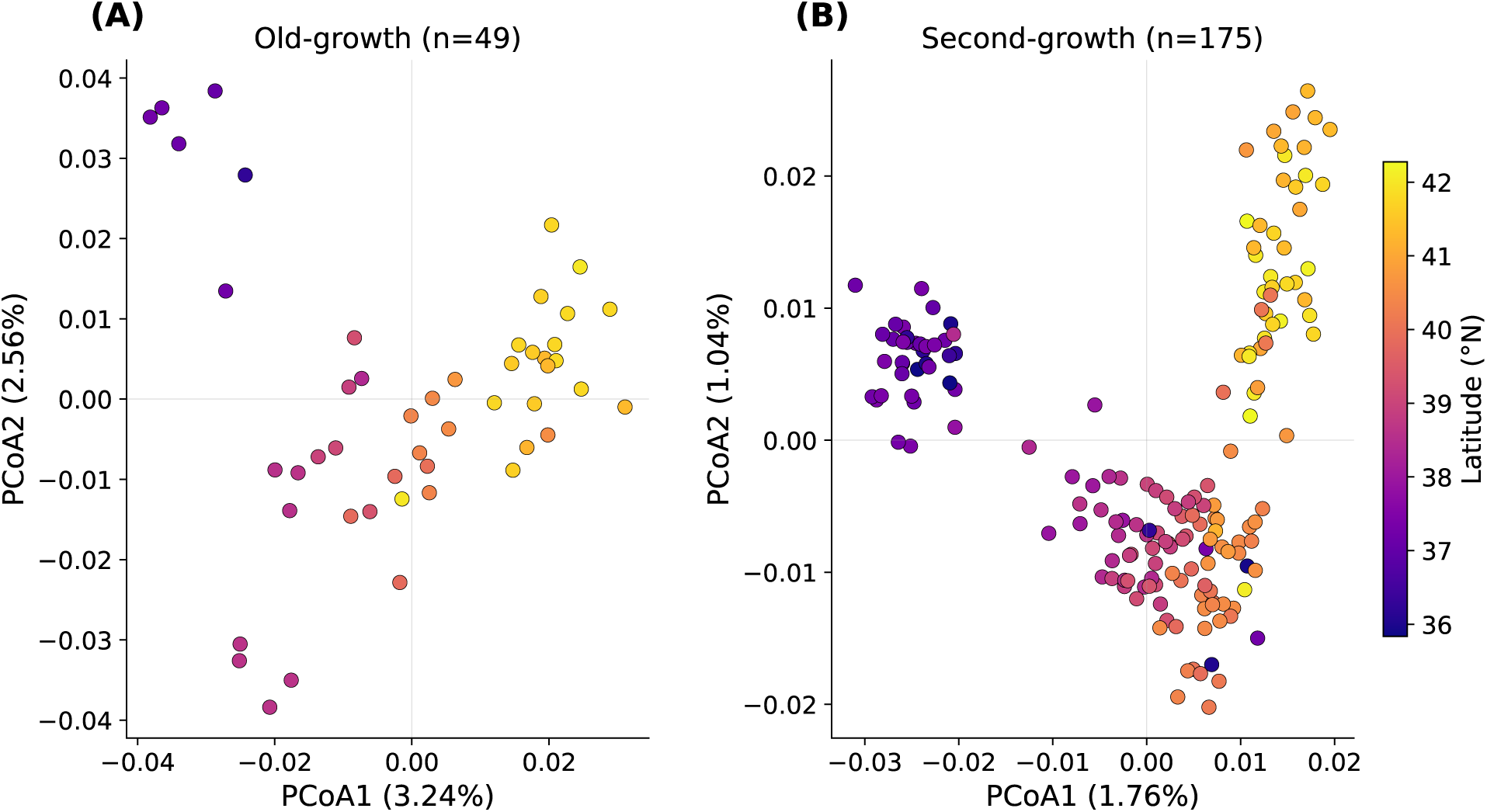
PCoA performed separately on **(A)** old-growth (*n* = 49) and **(B)** second-growth (*n* = 175) subsets of the 224-sample corrected IBD distance matrix. Points are coloured by latitude using a shared scale. The latitudinal gradient along PCoA1 is present in both subsets, confirming that the clinal structure is not an artefact of mixing forest types. The northern cluster is less pronounced in the old-growth subset, consistent with lower old-growth sampling density in the northern part of the range.

## Supplementary Materials

**Figure S1:**
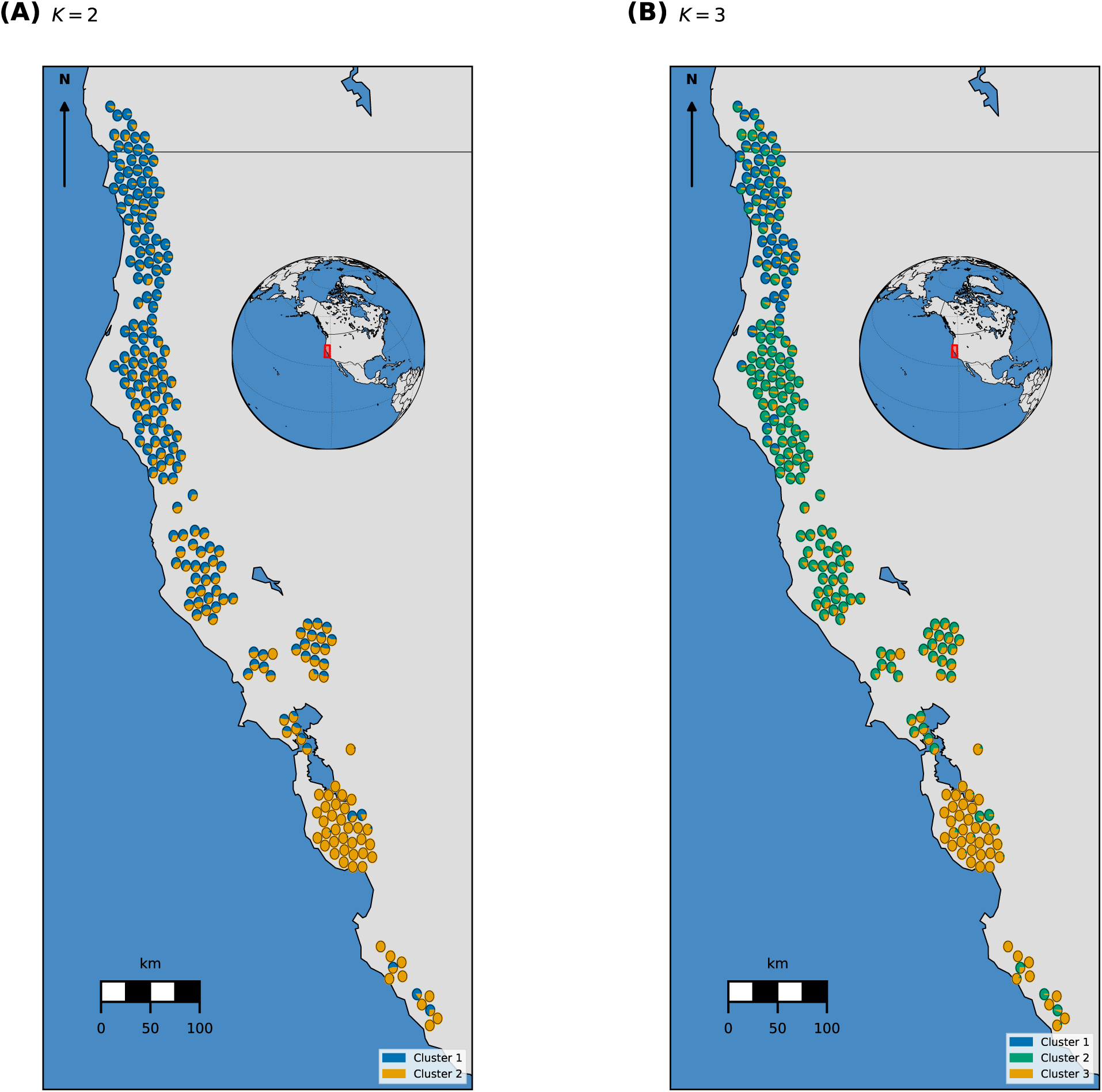
Geographic distribution of LEA sNMF admixture proportions. **(A)** *K* = 2; **(B)** *K* = 3. The southern genetic break occurs near the San Francisco Bay region (37.02°N) and the northern transition near the Humboldt–Mendocino county border (39.80°N), consistent with the segmented regression breakpoints.

Supplementary Table S1. (provided as a separate Excel file): IBD distances

Supplementary Table S2. (provided as a separate Excel file): top1000 fst.xlsx

Supplementary Table S3: Complete annotated list of *F_ST_* candidate loci (FST *×* LEA overlap SNPs and additional FST outliers on the same scaffolds).

**Table 2:** Complete list of annotated *F_ST_* candidate loci. **FST** *×* **LEA**: SNPs in both the top-1,000 *F_ST_* outliers and LFMM2 significant environmental associations (*q <* 0.05, K = 3). **FST outlier**: additional top-1,000 *F_ST_* outliers on the same redwood scaffolds as FST *×* LEA SNPs, directly annotated by BLASTx. All annotations against UniProt/SwissProt using diamond blastx (e-value *≤* 10*^−^*^4^, identity *≥* 35%). Env. variables: MAT = mean annual temperature, MTwQ = mean temp. of warmest quarter, MAP = mean annual precipitation, PDM = precip. driest month, P_season_ = precip. seasonality, PWQ = precip. warmest quarter, CEC = cation exchange capacity.

| Source | $F_{ST}$ | Gene | Putative function | Env. variables | SEGI position |
| --- | --- | --- | --- | --- | --- |
| **$F_{ST} \times LEA$ overlap SNPs (all 52)** |  |  |  |  |  |
| $F_{ST} \times LEA$ | 0.2511 | *MKK9* | Mitogen-activated protein kinase kinase 9 | MAP, MAT, MTwQ, $P_{season}$, PWQ | Chr11: 47.8 Mb |
| $F_{ST} \times LEA$ | 0.1139 | — | — | $P_{season}$, PDM, PWQ | Chr11: 86.3 Mb |
| $F_{ST} \times LEA$ | 0.1077 | *Fra* | Major strawberry allergen Fra a 1.07 | CEC, PDM, PWQ | Chr4: 578.0 Mb |
| $F_{ST} \times LEA$ | 0.1076 | — | — | $P_{season}$, PDM | Chr11: 2.7 Mb |
| $F_{ST} \times LEA$ | 0.1059 | — | — | $P_{season}$, PDM | Chr11: 2.7 Mb |
| $F_{ST} \times LEA$ | 0.1053 | — | — | $P_{season}$, PDM | Chr11: 2.7 Mb |
| $F_{ST} \times LEA$ | 0.1023 | *GYRB* | DNA gyrase subunit B, chloroplastic/mitochondrial | $P_{season}$ | Chr9: 475.1 Mb |

Table 2 – *continued*
| Source | $F_{ST}$ | Gene | Putative function | Env. variables | SEGI position |
| --- | --- | --- | --- | --- | --- |
| FST ×<br>LEA | 0.1006 | *RLK7* | Receptor-like protein kinase 7 | P<sub>season</sub>, PDM,<br>PWQ | Chr3:<br>259.5 Mb |
| FST ×<br>LEA | 0.0969 | *At5g48900* | Probable pectate lyase 20 | CEC, P<sub>season</sub>,<br>PDM, PWQ | Chr10:<br>102.3 Mb |
| FST ×<br>LEA | 0.0921 | *HSL1* | Receptor-like protein kinase<br>HSL1 | P<sub>season</sub>, PDM,<br>PWQ | Chr3:<br>259.5 Mb |
| FST ×<br>LEA | 0.0905 | *GSVIVT00020320001* | Probable polygalacturonase | PDM | Chr8:<br>63.7 Mb |
| FST ×<br>LEA | 0.0878 | *CEPR1* | Receptor protein-tyrosine kinase<br>CEPR1 | P<sub>season</sub>, PDM,<br>PWQ | Chr3:<br>259.5 Mb |
| FST ×<br>LEA | 0.0871 | *CEPR1* | Receptor protein-tyrosine kinase<br>CEPR1 | P<sub>season</sub>, PDM,<br>PWQ | Chr3:<br>259.5 Mb |
| FST ×<br>LEA | 0.0870 | — | — | PDM | Chr8:<br>87.5 Mb |
| FST ×<br>LEA | 0.0829 | *SWI3B* | SWI/SNF complex subunit<br>SWI3B | CEC | Chr4:<br>582.4 Mb |
| FST ×<br>LEA | 0.0781 | *TMN2* | Transmembrane 9 superfamily<br>member 2 | PDM | Chr8:<br>81.8 Mb |
| FST ×<br>LEA | 0.0778 | *OPR1* | 12-oxophytodienoate reductase 1 | P<sub>season</sub> | Chr3:<br>315.2 Mb |
| FST ×<br>LEA | 0.0764 | *DF1* | Trihelix transcription factor DF1 | PDM | Chr8: 5.7 Mb |
| FST ×<br>LEA | 0.0758 | *HOX11* | Homeobox-leucine zipper protein<br>HOX11 | PDM | Chr5:<br>243.5 Mb |
| FST ×<br>LEA | 0.0739 | — | — | PDM | Chr7:<br>17.8 Mb |

Table 2 – *continued*
| Source | $F_{ST}$ | Gene | Putative function | Env. variables | SEGI position |
| --- | --- | --- | --- | --- | --- |
| FST ×<br>LEA | 0.0734 | *SD25* | G-type lectin S-receptor-like<br>serine/threonine-protein kinase<br>SD2-5 | CEC | Chr7:<br>311.3 Mb |
| FST ×<br>LEA | 0.0732 | *At5g46100* | Pentatricopeptide<br>repeat-containing protein<br>At5g46100 | P<sub>season</sub>, PDM,<br>PWQ | Chr7:<br>79.4 Mb |
| FST ×<br>LEA | 0.0721 | — | — | P<sub>season</sub>, PDM,<br>PWQ | Chr7:<br>83.1 Mb |
| FST ×<br>LEA | 0.0719 | *NPF8.1* | Protein NRT1/ PTR FAMILY<br>8.1 | CEC, P<sub>season</sub>, PDM | Chr9:<br>114.9 Mb |
| FST ×<br>LEA | 0.0704 | *At3g02645* | Putative UPF0481 protein<br>At3g02645 | PDM | Chr1:<br>482.6 Mb |
| FST ×<br>LEA | 0.0690 | *Herc6* | E3 ISG15-protein ligase Herc6 | P<sub>season</sub>, PDM,<br>PWQ | Chr7:<br>620.4 Mb |
| FST ×<br>LEA | 0.0688 | — | — | PDM | Chr7:<br>83.1 Mb |
| FST ×<br>LEA | 0.0676 | *LAC3* | Laccase-3 | PDM | Chr4:<br>593.0 Mb |
| FST ×<br>LEA | 0.0675 | *At4g16563* | Probable aspartyl protease<br>At4g16563 | PDM, PWQ | Chr10:<br>612.6 Mb |
| FST ×<br>LEA | 0.0671 | — | — | P<sub>season</sub>, PDM,<br>PWQ | Chr1:<br>296.5 Mb |
| FST ×<br>LEA | 0.0667 | *psaA* | Photosystem I P700 chlorophyll a<br>apoprotein A1 | PDM | Chr7:<br>561.0 Mb |
| FST ×<br>LEA | 0.0666 | *LRK10* | Rust resistance kinase Lr10 | P<sub>season</sub>, PDM,<br>PWQ | Chr4:<br>101.3 Mb |

Table 2 – *continued*
| Source | $F_{ST}$ | Gene | Putative function | Env. variables | SEGI position |
| --- | --- | --- | --- | --- | --- |
| FST ×<br>LEA | 0.0665 | *XTH8* | Probable xyloglucan<br>endotransglucosylase/hydrolase<br>protein 8 | PDM | Chr1:<br>458.9 Mb |
| FST ×<br>LEA | 0.0660 | — | — | CEC | Chr4:<br>582.4 Mb |
| FST ×<br>LEA | 0.0652 | *RNR1* | Ribonucleoside-diphosphate<br>reductase large subunit | P<sub>season</sub>, PDM | Chr5:<br>588.5 Mb |
| FST ×<br>LEA | 0.0650 | *QRT3* | Polygalacturonase QRT3 | P<sub>season</sub>, PDM,<br>PWQ | Chr5:<br>120.1 Mb |
| FST ×<br>LEA | 0.0648 | *QRT3* | Polygalacturonase QRT3 | P<sub>season</sub>, PDM,<br>PWQ | Chr5:<br>120.1 Mb |
| FST ×<br>LEA | 0.0645 | *XTH8* | Probable xyloglucan<br>endotransglucosylase/hydrolase<br>protein 8 | PDM, PWQ | Chr1:<br>458.9 Mb |
| FST ×<br>LEA | 0.0637 | *HSP18.1* | 18.1 kDa class I heat shock<br>protein | P<sub>season</sub>, PDM | Chr3:<br>311.3 Mb |
| FST ×<br>LEA | 0.0636 | *SMC6B* | Structural maintenance of<br>chromosomes protein 6B | PDM | Chr4:<br>646.0 Mb |
| FST ×<br>LEA | 0.0634 | *TMN2* | Transmembrane 9 superfamily<br>member 2 | CEC, P<sub>season</sub>,<br>PDM, PWQ | Chr8:<br>81.8 Mb |
| FST ×<br>LEA | 0.0627 | *DRB6* | Double-stranded RNA-binding<br>protein 6 | P<sub>season</sub>, PDM | Chr8:<br>643.2 Mb |
| FST ×<br>LEA | 0.0618 | *Os04g0617900* | Germin-like protein 4-1 | P<sub>season</sub>, PDM,<br>PWQ | Chr3:<br>304.2 Mb |
| FST ×<br>LEA | 0.0617 | *THE1* | Receptor-like protein kinase<br>THESEUS 1 | PDM | Chr7:<br>193.6 Mb |

Table 2 – *continued*
| Source | $F_{ST}$ | Gene | Putative function | Env. variables | SEGI position |
| --- | --- | --- | --- | --- | --- |
| FST ×<br>LEA | 0.0611 | — | — | P<sub>season</sub>, PDM | Chr1:<br>127.9 Mb |
| FST ×<br>LEA | 0.0610 | *GSO2* | LRR receptor-like<br>serine/threonine-protein kinase<br>GSO2 | PDM, PWQ | Chr4:<br>121.3 Mb |
| FST ×<br>LEA | 0.0604 | *TY3B-I* | Transposon Ty3-I Gag-Pol<br>polyprotein | P<sub>season</sub>, PDM,<br>PWQ | Chr2:<br>114.1 Mb |
| FST ×<br>LEA | 0.0597 | — | — | PDM, PWQ | Chr8:<br>642.8 Mb |
| FST ×<br>LEA | 0.0587 | *P5CS* | Delta-1-pyrroline-5-carboxylate<br>synthase | CEC, P<sub>season</sub>,<br>PDM, PWQ | Chr9:<br>605.5 Mb |
| FST ×<br>LEA | 0.0585 | *NPF8.1* | Protein NRT1/ PTR FAMILY<br>8.1 | PDM | Chr9:<br>114.9 Mb |
| FST ×<br>LEA | 0.0580 | — | — | PDM, PWQ | Chr8:<br>642.8 Mb |
| FST ×<br>LEA | 0.0578 | — | — | PDM | Chr11:<br>49.1 Mb |
| **Additional FST outliers on same scaffolds (direct BLASTx)** |  |  |  |  |  |
| FST<br>outlier | 0.2234 | — | — | — | — |
| FST<br>outlier | 0.1848 | *Dctpp1* | dCTP pyrophosphatase 1 | — | — |
| FST<br>outlier | 0.1691 | *RBP47B* | Polyadenylate-binding protein<br>RBP47B | — | — |
| FST<br>outlier | 0.1614 | *LPR1* | Multicopper oxidase LPR1 | — | — |

Table 2 – *continued*
| Source | $F_{ST}$ | Gene | Putative function | Env. variables | SEGI position |
| --- | --- | --- | --- | --- | --- |
| FST outlier | 0.1565 | *CCOAOMT* | Caffeoyl-CoA<br>O-methyltransferase | — | — |
| FST outlier | 0.1350 | *VIII-A* | Myosin-3 | — | — |
| FST outlier | 0.1348 | *GAD1* | Glutamate decarboxylase 1 | — | — |
| FST outlier | 0.1341 | — | — | — | — |
| FST outlier | 0.1286 | *UGT89B1* | Flavonol 3-O-glucosyltransferase<br>UGT89B1 | — | — |
| FST outlier | 0.1279 | *OPR1* | 12-oxophytodienoate reductase 1 | — | — |
| FST outlier | 0.1229 | — | — | — | — |
| FST outlier | 0.1071 | *RBOHC* | Respiratory burst oxidase<br>homolog protein C | — | — |
| FST outlier | 0.1071 | *CDC20-2* | Cell division cycle 20.2, cofactor<br>of APC complex | — | — |
| FST outlier | 0.1039 | — | — | — | — |
| FST outlier | 0.1028 | — | — | — | — |
| FST outlier | 0.1024 | *daf-2* | Insulin-like receptor | — | — |
| FST outlier | 0.1022 | — | — | — | — |

Table 2 – *continued*
| Source | $F_{ST}$ | Gene | Putative function | Env. variables | SEGI position |
| --- | --- | --- | --- | --- | --- |
| FST outlier | 0.0948 | — | — | — | — |
| FST outlier | 0.0848 | *NAGK* | Acetylglutamate kinase, chloroplastic | — | — |
| FST outlier | 0.0780 | *F3H-2* | Flavanone 3-dioxygenase 2 | — | — |

Supplementary Table S4: Co-localized *F_ST_* and LFMM2 genomic windows (non-exact overlap; 1-Mb co-localization windows).

**Table S4:** Co-localized *F_ST_* and LFMM2 genomic windows. Each row represents a 1-Mb window on the giant sequoia (*SEGI* 2.0) reference genome where at least one *F_ST_*outlier SNP (top 1,000) and at least one LFMM2-significant SNP (*q <* 0.05) are present within 1 Mb but do not share the same SNP identifier. *n*_FST_ and *n*_LEA_: number of unique SNPs from each analysis in the window. Top *F_ST_* : highest *F_ST_* value among FST SNPs in the window, with the containing scaffold. Top *−* log_10_(*p*): strongest LFMM2 signal in the window. Dominant env. variable: most frequent LFMM2 association variable among LEA SNPs in the window.

| Chr | Region (Mb) | $n_{FST}$ | $n_{LEA}$ | Top $F_{ST}$ | Top $-\log_{10}(p)$ | Top FST scaffold | Dominant env. variable |
| --- | --- | --- | --- | --- | --- | --- | --- |
| chr1 | 18–19 | 2 | 1 | 0.064 | 4.15 | HRSCAF=155548 | bio14 (precip driest month) |
| chr1 | 77–78 | 3 | 2 | 0.079 | 4.92 | HRSCAF=367399 | bio10 (mean temp warmest qtr) |
| chr1 | 154–155 | 1 | 1 | 0.107 | 4.66 | HRSCAF=412686 | bio4 (temp seasonality) |
| chr1 | 166–167 | 1 | 2 | 0.087 | 4.79 | HRSCAF=166073 | bio14 (precip driest month) |
| chr1 | 175–176 | 1 | 3 | 0.059 | 5.69 | HRSCAF=196999 | CEC 0–5 cm |
| chr1 | 193–194 | 1 | 2 | 0.059 | 5.51 | HRSCAF=166073 | bio10 (mean temp warmest qtr) |
| chr1 | 198–199 | 1 | 1 | 0.057 | 6.65 | HRSCAF=219368 | bio14 (precip driest month) |
| chr1 | 258–259 | 3 | 3 | 0.104 | 5.17 | HRSCAF=182310 | bio14 (precip driest month) |
| chr1 | 274–275 | 1 | 4 | 0.062 | 8.26 | HRSCAF=7118 | bio14 (precip driest month) |

Table S4 (continued)
| Chr | Region (Mb) | $n_{\text{FST}}$ | $n_{\text{LEA}}$ | Top $F_{ST}$ | Top $-\log_{10}(p)$ | Top FST scaffold | Dominant env. variable |
| --- | --- | --- | --- | --- | --- | --- | --- |
| chr1 | 299–300 | 1 | 1 | 0.062 | 5.06 | HRSCAF=7341 | bio14 (precip driest month) |
| chr1 | 313–314 | 1 | 1 | 0.065 | 5.09 | HRSCAF=7341 | bio14 (precip driest month) |
| chr1 | 452–453 | 1 | 1 | 0.058 | 5.98 | HRSCAF=347886 | bio4 (temp seasonality) |
| chr1 | 458–459 | 5 | 6 | 0.081 | 7.73 | HRSCAF=367399 | bio14 (precip driest month) |
| chr1 | 845–846 | 1 | 1 | 0.072 | 4.24 | HRSCAF=412686 | bio10 (mean temp warmest qtr) |
| chr1 | 884–885 | 1 | 1 | 0.107 | 4.87 | HRSCAF=211191 | bio5 (max temp warmest month) |
| chr1 | 936–937 | 2 | 1 | 0.068 | 6.53 | HRSCAF=347886 | bio14 (precip driest month) |
| chr1 | 952–953 | 1 | 3 | 0.101 | 4.80 | HRSCAF=155469 | bio14 (precip driest month) |
| chr1 | 971–972 | 1 | 1 | 0.065 | 5.73 | HRSCAF=155548 | bio10 (mean temp warmest qtr) |
| chr1 | 972–973 | 1 | 1 | 0.058 | 5.73 | HRSCAF=155548 | bio10 (mean temp warmest qtr) |
| chr2 | 13–14 | 1 | 3 | 0.088 | 5.94 | HRSCAF=166073 | bio14 (precip driest month) |
| chr2 | 35–36 | 1 | 1 | 0.062 | 6.18 | HRSCAF=293701 | bio1 (annual mean temp) |
| chr2 | 95–96 | 1 | 1 | 0.063 | 7.09 | HRSCAF=7118 | bio10 (mean temp warmest qtr) |
| chr2 | 114–115 | 1 | 1 | 0.071 | 4.82 | HRSCAF=7118 | bio14 (precip driest month) |
| chr2 | 142–143 | 1 | 1 | 0.057 | 4.20 | HRSCAF=7118 | bio14 (precip driest month) |
| chr2 | 218–219 | 1 | 1 | 0.070 | 4.51 | HRSCAF=166073 | bio14 (precip driest month) |
| chr2 | 241–242 | 1 | 1 | 0.058 | 4.23 | HRSCAF=166073 | bio14 (precip driest month) |
| chr2 | 292–293 | 2 | 1 | 0.062 | 4.34 | HRSCAF=155548 | bio14 (precip driest month) |
| chr2 | 303–304 | 1 | 1 | 0.061 | 3.95 | HRSCAF=366706 | bio14 (precip driest month) |
| chr2 | 382–383 | 4 | 1 | 0.075 | 4.22 | HRSCAF=7118 | bio10 (mean temp warmest qtr) |
| chr2 | 396–397 | 1 | 1 | 0.060 | 5.21 | HRSCAF=219368 | bio10 (mean temp warmest qtr) |
| chr2 | 522–523 | 2 | 1 | 0.061 | 5.67 | HRSCAF=7341 | bio10 (mean temp warmest qtr) |
| chr2 | 573–574 | 1 | 1 | 0.062 | 4.08 | HRSCAF=377209 | bio14 (precip driest month) |
| chr2 | 620–621 | 1 | 6 | 0.069 | 7.60 | HRSCAF=377209 | bio14 (precip driest month) |

Table S4 (continued)
| Chr | Region (Mb) | $n_{\text{FST}}$ | $n_{\text{LEA}}$ | Top $F_{ST}$ | Top $-\log_{10}(p)$ | Top FST scaffold | Dominant env. variable |
| --- | --- | --- | --- | --- | --- | --- | --- |
| chr2 | 632–633 | 1 | 1 | 0.064 | 4.30 | HRSCAF=219368 | bio14 (precip driest month) |
| chr2 | 693–694 | 4 | 2 | 0.128 | 5.26 | HRSCAF=7118 | bio10 (mean temp warmest qtr) |
| chr2 | 713–714 | 8 | 3 | 0.074 | 6.99 | HRSCAF=219368 | bio14 (precip driest month) |
| chr2 | 744–745 | 1 | 3 | 0.063 | 5.51 | HRSCAF=7341 | bio4 (temp seasonality) |
| chr2 | 745–746 | 1 | 3 | 0.061 | 5.51 | HRSCAF=7341 | bio4 (temp seasonality) |
| chr2 | 757–758 | 1 | 1 | 0.088 | 4.78 | HRSCAF=412686 | bio10 (mean temp warmest qtr) |
| chr2 | 759–760 | 1 | 1 | 0.068 | 4.61 | HRSCAF=7341 | bio10 (mean temp warmest qtr) |
| chr2 | 768–769 | 1 | 1 | 0.090 | 4.39 | HRSCAF=7341 | bio14 (precip driest month) |
| chr2 | 769–770 | 2 | 1 | 0.116 | 4.39 | HRSCAF=7341 | bio14 (precip driest month) |
| chr2 | 773–774 | 3 | 2 | 0.077 | 7.66 | HRSCAF=219368 | bio5 (max temp warmest month) |
| chr2 | 808–809 | 1 | 2 | 0.059 | 4.98 | HRSCAF=29663 | bio14 (precip driest month) |
| chr2 | 809–810 | 2 | 2 | 0.093 | 4.98 | HRSCAF=29663 | bio14 (precip driest month) |
| chr2 | 834–835 | 2 | 1 | 0.061 | 5.00 | HRSCAF=367399 | bio14 (precip driest month) |
| chr2 | 851–852 | 3 | 1 | 0.075 | 3.99 | HRSCAF=293701 | bio14 (precip driest month) |
| chr2 | 867–868 | 1 | 2 | 0.082 | 4.20 | HRSCAF=293701 | bio14 (precip driest month) |
| chr3 | 0–1 | 1 | 5 | 0.068 | 5.08 | HRSCAF=367399 | bio10 (mean temp warmest qtr) |
| chr3 | 5–6 | 1 | 1 | 0.087 | 3.99 | HRSCAF=412686 | bio14 (precip driest month) |
| chr3 | 40–41 | 1 | 1 | 0.065 | 4.48 | HRSCAF=7341 | bio14 (precip driest month) |
| chr3 | 42–43 | 1 | 1 | 0.059 | 6.51 | HRSCAF=412686 | bio10 (mean temp warmest qtr) |
| chr3 | 43–44 | 1 | 1 | 0.068 | 6.51 | HRSCAF=412686 | bio10 (mean temp warmest qtr) |
| chr3 | 78–79 | 1 | 1 | 0.062 | 4.46 | HRSCAF=367399 | bio14 (precip driest month) |
| chr3 | 110–111 | 7 | 1 | 0.069 | 4.58 | HRSCAF=182310 | bio10 (mean temp warmest qtr) |
| chr3 | 188–189 | 1 | 1 | 0.065 | 8.01 | HRSCAF=211191 | bio10 (mean temp warmest qtr) |
| chr3 | 196–197 | 1 | 1 | 0.073 | 4.75 | HRSCAF=412686 | bio4 (temp seasonality) |

Table S4 (continued)
| Chr | Region (Mb) | $n_{\text{FST}}$ | $n_{\text{LEA}}$ | Top $F_{ST}$ | Top $-\log_{10}(p)$ | Top FST scaffold | Dominant env. variable |
| --- | --- | --- | --- | --- | --- | --- | --- |
| chr3 | 230–231 | 2 | 2 | 0.063 | 5.84 | HRSCAF=166073 | bio14 (precip driest month) |
| chr3 | 239–240 | 3 | 2 | 0.070 | 4.57 | HRSCAF=366706 | bio14 (precip driest month) |
| chr3 | 246–247 | 1 | 1 | 0.063 | 4.44 | HRSCAF=415217 | bio10 (mean temp warmest qtr) |
| chr3 | 253–254 | 2 | 1 | 0.064 | 4.94 | HRSCAF=367399 | bio14 (precip driest month) |
| chr3 | 274–275 | 1 | 1 | 0.089 | 4.04 | HRSCAF=366706 | bio14 (precip driest month) |
| chr3 | 284–285 | 1 | 1 | 0.082 | 4.04 | HRSCAF=366706 | bio14 (precip driest month) |
| chr3 | 285–286 | 1 | 1 | 0.075 | 4.04 | HRSCAF=366706 | bio14 (precip driest month) |
| chr3 | 311–312 | 6 | 1 | 0.097 | 4.55 | HRSCAF=196999 | bio14 (precip driest month) |
| chr3 | 328–329 | 1 | 1 | 0.067 | 3.98 | HRSCAF=293701 | bio14 (precip driest month) |
| chr3 | 329–330 | 1 | 1 | 0.091 | 3.98 | HRSCAF=367399 | bio14 (precip driest month) |
| chr3 | 510–511 | 1 | 1 | 0.093 | 5.06 | HRSCAF=377209 | bio10 (mean temp warmest qtr) |
| chr3 | 683–684 | 2 | 3 | 0.074 | 4.18 | HRSCAF=219368 | bio14 (precip driest month) |
| chr3 | 686–687 | 1 | 2 | 0.068 | 4.72 | HRSCAF=219368 | bio10 (mean temp warmest qtr) |
| chr3 | 690–691 | 5 | 1 | 0.092 | 4.71 | HRSCAF=219368 | bio14 (precip driest month) |
| chr3 | 753–754 | 1 | 1 | 0.074 | 4.98 | HRSCAF=377209 | bio14 (precip driest month) |
| chr3 | 816–817 | 1 | 1 | 0.057 | 4.71 | HRSCAF=7341 | bio14 (precip driest month) |
| chr3 | 817–818 | 1 | 1 | 0.068 | 4.05 | HRSCAF=192499 | bio14 (precip driest month) |
| chr4 | 14–15 | 1 | 1 | 0.057 | 4.69 | HRSCAF=90849 | bio4 (temp seasonality) |
| chr4 | 101–102 | 1 | 2 | 0.063 | 4.90 | HRSCAF=377209 | bio14 (precip driest month) |
| chr4 | 120–121 | 1 | 4 | 0.064 | 6.33 | HRSCAF=377209 | bio14 (precip driest month) |
| chr4 | 121–122 | 6 | 6 | 0.072 | 8.04 | HRSCAF=377209 | bio14 (precip driest month) |
| chr4 | 184–185 | 3 | 2 | 0.104 | 4.30 | HRSCAF=293701 | bio10 (mean temp warmest qtr) |
| chr4 | 185–186 | 4 | 1 | 0.126 | 4.02 | HRSCAF=293701 | bio14 (precip driest month) |
| chr4 | 249–250 | 1 | 1 | 0.078 | 4.35 | HRSCAF=90849 | bio14 (precip driest month) |

Table S4 (continued)
| Chr | Region (Mb) | $n_{\text{FST}}$ | $n_{\text{LEA}}$ | Top $F_{ST}$ | Top $-\log_{10}(p)$ | Top FST scaffold | Dominant env. variable |
| --- | --- | --- | --- | --- | --- | --- | --- |
| chr4 | 308–309 | 1 | 3 | 0.065 | 5.23 | HRSCAF=155548 | bio14 (precip driest month) |
| chr4 | 504–505 | 2 | 3 | 0.070 | 5.55 | HRSCAF=219368 | bio14 (precip driest month) |
| chr4 | 545–546 | 2 | 2 | 0.068 | 5.02 | HRSCAF=377209 | bio14 (precip driest month) |
| chr4 | 581–582 | 2 | 6 | 0.090 | 6.54 | HRSCAF=219368 | bio10 (mean temp warmest qtr) |
| chr4 | 582–583 | 1 | 5 | 0.072 | 6.54 | HRSCAF=219368 | bio10 (mean temp warmest qtr) |
| chr4 | 600–601 | 7 | 2 | 0.073 | 4.54 | HRSCAF=155548 | bio10 (mean temp warmest qtr) |
| chr4 | 645–646 | 1 | 3 | 0.071 | 6.32 | HRSCAF=367399 | bio5 (max temp warmest month) |
| chr4 | 647–648 | 1 | 1 | 0.063 | 5.81 | HRSCAF=17205 | CEC 0–5 cm |
| chr4 | 656–657 | 1 | 1 | 0.057 | 5.27 | HRSCAF=351831 | bio10 (mean temp warmest qtr) |
| chr4 | 677–678 | 1 | 1 | 0.063 | 5.43 | HRSCAF=377209 | bio14 (precip driest month) |
| chr4 | 683–684 | 1 | 1 | 0.062 | 4.68 | HRSCAF=377209 | bio4 (temp seasonality) |
| chr5 | 99–100 | 1 | 1 | 0.060 | 4.92 | HRSCAF=182310 | bio10 (mean temp warmest qtr) |
| chr5 | 103–104 | 2 | 2 | 0.092 | 5.98 | HRSCAF=96993 | bio10 (mean temp warmest qtr) |
| chr5 | 199–200 | 1 | 1 | 0.068 | 3.94 | HRSCAF=412686 | bio14 (precip driest month) |
| chr5 | 230–231 | 1 | 3 | 0.092 | 7.07 | HRSCAF=412686 | bio10 (mean temp warmest qtr) |
| chr5 | 231–232 | 1 | 3 | 0.058 | 7.07 | HRSCAF=412686 | bio10 (mean temp warmest qtr) |
| chr5 | 266–267 | 5 | 1 | 0.087 | 5.14 | HRSCAF=377209 | bio14 (precip driest month) |
| chr5 | 377–378 | 1 | 2 | 0.059 | 4.91 | HRSCAF=367399 | bio14 (precip driest month) |
| chr5 | 545–546 | 1 | 2 | 0.060 | 5.22 | HRSCAF=366706 | bio14 (precip driest month) |
| chr5 | 546–547 | 1 | 2 | 0.061 | 5.22 | HRSCAF=366706 | bio14 (precip driest month) |
| chr5 | 565–566 | 4 | 1 | 0.108 | 4.86 | HRSCAF=219368 | bio10 (mean temp warmest qtr) |
| chr6 | 1–2 | 1 | 3 | 0.062 | 5.93 | HRSCAF=366706 | bio10 (mean temp warmest qtr) |
| chr6 | 26–27 | 1 | 1 | 0.071 | 4.31 | HRSCAF=366706 | bio10 (mean temp warmest qtr) |
| chr6 | 48–49 | 1 | 1 | 0.060 | 5.77 | HRSCAF=366706 | bio10 (mean temp warmest qtr) |

Table S4 (continued)
| Chr | Region (Mb) | $n_{\text{FST}}$ | $n_{\text{LEA}}$ | Top $F_{ST}$ | Top $-\log_{10}(p)$ | Top FST scaffold | Dominant env. variable |
| --- | --- | --- | --- | --- | --- | --- | --- |
| chr6 | 64–65 | 1 | 4 | 0.086 | 7.50 | HRSCAF=90849 | bio14 (precip driest month) |
| chr6 | 79–80 | 1 | 2 | 0.069 | 5.11 | HRSCAF=155548 | bio14 (precip driest month) |
| chr6 | 101–102 | 1 | 1 | 0.058 | 4.93 | HRSCAF=377209 | bio14 (precip driest month) |
| chr6 | 115–116 | 1 | 3 | 0.100 | 4.80 | HRSCAF=155548 | bio14 (precip driest month) |
| chr6 | 162–163 | 1 | 1 | 0.061 | 5.36 | HRSCAF=75987 | bio10 (mean temp warmest qtr) |
| chr6 | 183–184 | 1 | 3 | 0.060 | 4.98 | HRSCAF=90849 | bio10 (mean temp warmest qtr) |
| chr6 | 184–185 | 2 | 2 | 0.069 | 5.58 | HRSCAF=90849 | bio10 (mean temp warmest qtr) |
| chr6 | 185–186 | 1 | 2 | 0.058 | 5.58 | HRSCAF=182310 | bio10 (mean temp warmest qtr) |
| chr6 | 202–203 | 1 | 1 | 0.067 | 5.47 | HRSCAF=367399 | bio10 (mean temp warmest qtr) |
| chr6 | 206–207 | 1 | 3 | 0.071 | 4.87 | HRSCAF=182310 | bio10 (mean temp warmest qtr) |
| chr6 | 460–461 | 1 | 1 | 0.074 | 5.85 | HRSCAF=219368 | bio15 (precip seasonality) |
| chr6 | 476–477 | 1 | 1 | 0.072 | 4.07 | HRSCAF=412686 | bio14 (precip driest month) |
| chr6 | 503–504 | 1 | 1 | 0.060 | 5.10 | HRSCAF=219368 | bio10 (mean temp warmest qtr) |
| chr6 | 504–505 | 3 | 1 | 0.069 | 5.10 | HRSCAF=96993 | bio10 (mean temp warmest qtr) |
| chr6 | 516–517 | 2 | 1 | 0.084 | 6.01 | HRSCAF=211191 | bio10 (mean temp warmest qtr) |
| chr6 | 540–541 | 2 | 1 | 0.087 | 4.06 | HRSCAF=110562 | bio14 (precip driest month) |
| chr6 | 627–628 | 1 | 2 | 0.063 | 5.68 | HRSCAF=366706 | bio14 (precip driest month) |
| chr6 | 642–643 | 1 | 4 | 0.067 | 4.71 | HRSCAF=366706 | bio14 (precip driest month) |
| chr6 | 655–656 | 1 | 2 | 0.076 | 5.36 | HRSCAF=41004 | bio14 (precip driest month) |
| chr7 | 27–28 | 3 | 1 | 0.064 | 5.08 | HRSCAF=17205 | bio10 (mean temp warmest qtr) |
| chr7 | 58–59 | 2 | 2 | 0.062 | 3.95 | HRSCAF=366706 | bio14 (precip driest month) |
| chr7 | 86–87 | 3 | 2 | 0.114 | 5.98 | HRSCAF=155707 | bio14 (precip driest month) |
| chr7 | 89–90 | 1 | 2 | 0.089 | 5.14 | HRSCAF=155707 | bio18 (precip warmest qtr) |
| chr7 | 90–91 | 1 | 3 | 0.060 | 5.49 | HRSCAF=90849 | bio18 (precip warmest qtr) |

Table S4 (continued)
| Chr | Region (Mb) | $n_{\text{FST}}$ | $n_{\text{LEA}}$ | Top $F_{ST}$ | Top $-\log_{10}(p)$ | Top FST scaffold | Dominant env. variable |
| --- | --- | --- | --- | --- | --- | --- | --- |
| chr7 | 91–92 | 1 | 3 | 0.065 | 5.49 | HRSCAF=155707 | bio18 (precip warmest qtr) |
| chr7 | 111–112 | 2 | 1 | 0.094 | 4.66 | HRSCAF=155707 | bio14 (precip driest month) |
| chr7 | 130–131 | 3 | 2 | 0.104 | 5.71 | HRSCAF=166073 | CEC 0–5 cm |
| chr7 | 152–153 | 1 | 1 | 0.059 | 4.11 | HRSCAF=293701 | bio14 (precip driest month) |
| chr7 | 153–154 | 1 | 3 | 0.058 | 4.48 | HRSCAF=293701 | bio14 (precip driest month) |
| chr7 | 176–177 | 2 | 1 | 0.145 | 4.58 | HRSCAF=412686 | bio10 (mean temp warmest qtr) |
| chr7 | 189–190 | 1 | 3 | 0.065 | 7.17 | HRSCAF=366706 | bio14 (precip driest month) |
| chr7 | 300–301 | 1 | 1 | 0.071 | 4.82 | HRSCAF=166073 | bio10 (mean temp warmest qtr) |
| chr7 | 301–302 | 1 | 1 | 0.065 | 4.82 | HRSCAF=155548 | bio10 (mean temp warmest qtr) |
| chr7 | 302–303 | 1 | 1 | 0.061 | 4.82 | HRSCAF=347886 | bio10 (mean temp warmest qtr) |
| chr7 | 311–312 | 2 | 1 | 0.079 | 4.50 | HRSCAF=155548 | bio10 (mean temp warmest qtr) |
| chr7 | 542–543 | 2 | 1 | 0.101 | 4.02 | HRSCAF=96993 | bio14 (precip driest month) |
| chr7 | 553–554 | 2 | 1 | 0.069 | 8.59 | HRSCAF=367399 | bio10 (mean temp warmest qtr) |
| chr7 | 555–556 | 3 | 1 | 0.074 | 3.93 | HRSCAF=269722 | bio14 (precip driest month) |
| chr7 | 556–557 | 1 | 1 | 0.058 | 3.93 | HRSCAF=269722 | bio14 (precip driest month) |
| chr7 | 623–624 | 2 | 2 | 0.070 | 5.63 | HRSCAF=7341 | bio14 (precip driest month) |
| chr7 | 627–628 | 1 | 1 | 0.068 | 4.82 | HRSCAF=183235 | bio14 (precip driest month) |
| chr7 | 629–630 | 8 | 4 | 0.108 | 4.60 | HRSCAF=7341 | bio14 (precip driest month) |
| chr7 | 651–652 | 1 | 3 | 0.069 | 4.13 | HRSCAF=7341 | bio14 (precip driest month) |
| chr8 | 15–16 | 1 | 1 | 0.100 | 4.29 | HRSCAF=211191 | bio10 (mean temp warmest qtr) |
| chr8 | 23–24 | 1 | 2 | 0.068 | 4.74 | HRSCAF=367399 | bio14 (precip driest month) |
| chr8 | 63–64 | 2 | 2 | 0.072 | 4.46 | HRSCAF=412686 | bio14 (precip driest month) |
| chr8 | 76–77 | 1 | 3 | 0.092 | 6.82 | HRSCAF=219368 | bio14 (precip driest month) |
| chr8 | 89–90 | 3 | 6 | 0.082 | 5.16 | HRSCAF=219368 | bio15 (precip seasonality) |

Table S4 (continued)
| Chr | Region (Mb) | $n_{\text{FST}}$ | $n_{\text{LEA}}$ | Top $F_{ST}$ | Top $-\log_{10}(p)$ | Top FST scaffold | Dominant env. variable |
| --- | --- | --- | --- | --- | --- | --- | --- |
| chr8 | 98–99 | 2 | 3 | 0.084 | 5.70 | HRSCAF=377209 | bio10 (mean temp warmest qtr) |
| chr8 | 114–115 | 2 | 1 | 0.090 | 4.41 | HRSCAF=219368 | bio10 (mean temp warmest qtr) |
| chr8 | 133–134 | 1 | 2 | 0.069 | 5.72 | HRSCAF=182310 | bio14 (precip driest month) |
| chr8 | 159–160 | 1 | 1 | 0.058 | 4.95 | HRSCAF=110562 | bio10 (mean temp warmest qtr) |
| chr8 | 172–173 | 1 | 1 | 0.072 | 4.71 | HRSCAF=366706 | bio10 (mean temp warmest qtr) |
| chr8 | 201–202 | 1 | 1 | 0.064 | 4.81 | HRSCAF=377209 | bio10 (mean temp warmest qtr) |
| chr8 | 300–301 | 1 | 1 | 0.058 | 4.77 | HRSCAF=219368 | bio14 (precip driest month) |
| chr8 | 404–405 | 3 | 1 | 0.117 | 5.57 | HRSCAF=219368 | bio10 (mean temp warmest qtr) |
| chr8 | 501–502 | 1 | 1 | 0.057 | 4.09 | HRSCAF=293701 | bio14 (precip driest month) |
| chr8 | 642–643 | 1 | 3 | 0.085 | 6.49 | HRSCAF=367399 | bio14 (precip driest month) |
| chr9 | 27–28 | 1 | 1 | 0.078 | 4.02 | HRSCAF=347886 | bio14 (precip driest month) |
| chr9 | 36–37 | 2 | 3 | 0.084 | 5.91 | HRSCAF=219368 | bio14 (precip driest month) |
| chr9 | 53–54 | 1 | 2 | 0.065 | 5.88 | HRSCAF=367399 | bio14 (precip driest month) |
| chr9 | 69–70 | 1 | 3 | 0.066 | 6.67 | HRSCAF=90849 | bio14 (precip driest month) |
| chr9 | 84–85 | 3 | 2 | 0.099 | 4.20 | HRSCAF=377209 | bio14 (precip driest month) |
| chr9 | 110–111 | 2 | 2 | 0.088 | 5.82 | HRSCAF=377209 | bio14 (precip driest month) |
| chr9 | 121–122 | 1 | 7 | 0.091 | 6.85 | HRSCAF=412686 | bio14 (precip driest month) |
| chr9 | 122–123 | 3 | 4 | 0.112 | 5.21 | HRSCAF=412686 | bio14 (precip driest month) |
| chr9 | 136–137 | 2 | 1 | 0.060 | 5.18 | HRSCAF=377209 | bio10 (mean temp warmest qtr) |
| chr9 | 137–138 | 1 | 1 | 0.058 | 5.18 | HRSCAF=75987 | bio10 (mean temp warmest qtr) |
| chr9 | 146–147 | 1 | 4 | 0.058 | 6.66 | HRSCAF=412686 | bio14 (precip driest month) |
| chr9 | 517–518 | 2 | 1 | 0.064 | 4.09 | HRSCAF=18833 | bio14 (precip driest month) |
| chr9 | 572–573 | 2 | 4 | 0.067 | 5.40 | HRSCAF=293701 | bio10 (mean temp warmest qtr) |
| chr9 | 598–599 | 4 | 5 | 0.142 | 6.77 | HRSCAF=293701 | bio14 (precip driest month) |

Table S4 (continued)
| Chr | Region (Mb) | $n_{\text{FST}}$ | $n_{\text{LEA}}$ | Top $F_{ST}$ | Top $-\log_{10}(p)$ | Top FST scaffold | Dominant env. variable |
| --- | --- | --- | --- | --- | --- | --- | --- |
| chr9 | 609–610 | 1 | 2 | 0.064 | 5.35 | HRSCAF=366706 | bio14 (precip driest month) |
| chr9 | 631–632 | 1 | 1 | 0.072 | 5.04 | HRSCAF=7341 | bio10 (mean temp warmest qtr) |
| chr9 | 638–639 | 1 | 2 | 0.129 | 4.26 | HRSCAF=7341 | bio14 (precip driest month) |
| chr10 | 8–9 | 1 | 5 | 0.070 | 6.46 | HRSCAF=90849 | bio14 (precip driest month) |
| chr10 | 15–16 | 2 | 1 | 0.085 | 4.78 | HRSCAF=367399 | bio10 (mean temp warmest qtr) |
| chr10 | 26–27 | 2 | 1 | 0.102 | 4.86 | HRSCAF=367399 | bio10 (mean temp warmest qtr) |
| chr10 | 52–53 | 2 | 1 | 0.061 | 4.58 | HRSCAF=219368 | bio10 (mean temp warmest qtr) |
| chr10 | 98–99 | 1 | 1 | 0.073 | 4.98 | HRSCAF=29663 | bio14 (precip driest month) |
| chr10 | 106–107 | 1 | 3 | 0.075 | 8.91 | HRSCAF=90849 | bio14 (precip driest month) |
| chr10 | 111–112 | 1 | 1 | 0.057 | 6.55 | HRSCAF=90849 | bio10 (mean temp warmest qtr) |
| chr10 | 140–141 | 3 | 1 | 0.101 | 4.61 | HRSCAF=90849 | bio10 (mean temp warmest qtr) |
| chr10 | 528–529 | 2 | 1 | 0.089 | 4.67 | HRSCAF=135539 | bio10 (mean temp warmest qtr) |
| chr10 | 534–535 | 2 | 1 | 0.058 | 5.07 | HRSCAF=135539 | bio14 (precip driest month) |
| chr10 | 538–539 | 1 | 1 | 0.086 | 4.64 | HRSCAF=135539 | bio14 (precip driest month) |
| chr10 | 549–550 | 1 | 3 | 0.081 | 5.72 | HRSCAF=219368 | bio14 (precip driest month) |
| chr10 | 550–551 | 1 | 3 | 0.064 | 5.72 | HRSCAF=135539 | bio14 (precip driest month) |
| chr10 | 569–570 | 1 | 1 | 0.072 | 4.38 | HRSCAF=155548 | bio10 (mean temp warmest qtr) |
| chr10 | 570–571 | 2 | 3 | 0.080 | 5.40 | HRSCAF=155548 | bio14 (precip driest month) |
| chr10 | 577–578 | 1 | 2 | 0.060 | 4.72 | HRSCAF=155548 | bio14 (precip driest month) |
| chr10 | 609–610 | 7 | 5 | 0.080 | 5.10 | HRSCAF=90849 | bio14 (precip driest month) |
| chr10 | 612–613 | 1 | 2 | 0.074 | 5.79 | HRSCAF=7118 | bio14 (precip driest month) |
| chr10 | 627–628 | 1 | 1 | 0.069 | 4.90 | HRSCAF=293701 | bio14 (precip driest month) |
| chr11 | 46–47 | 2 | 1 | 0.112 | 5.52 | HRSCAF=412686 | CEC 0–5 cm |
| chr11 | 47–48 | 2 | 1 | 0.108 | 5.52 | HRSCAF=412686 | CEC 0–5 cm |

Table S4 (continued)
| Chr | Region (Mb) | $n_{\text{FST}}$ | $n_{\text{LEA}}$ | Top $F_{ST}$ | Top $-\log_{10}(p)$ | Top FST scaffold | Dominant env. variable |
| --- | --- | --- | --- | --- | --- | --- | --- |
| chr11 | 49–50 | 1 | 8 | 0.075 | 7.43 | HRSCAF=219368 | bio14 (precip driest month) |
| chr11 | 52–53 | 3 | 2 | 0.131 | 4.52 | HRSCAF=211191 | bio10 (mean temp warmest qtr) |
| chr11 | 106–107 | 2 | 1 | 0.077 | 4.54 | HRSCAF=219368 | bio14 (precip driest month) |
| chr11 | 127–128 | 2 | 1 | 0.062 | 3.95 | HRSCAF=351831 | bio14 (precip driest month) |
| chr11 | 262–263 | 2 | 3 | 0.064 | 5.77 | HRSCAF=166073 | bio10 (mean temp warmest qtr) |
| chr11 | 274–275 | 3 | 3 | 0.100 | 6.97 | HRSCAF=377209 | bio10 (mean temp warmest qtr) |
| chr11 | 335–336 | 1 | 2 | 0.076 | 6.24 | HRSCAF=17205 | bio14 (precip driest month) |
| chr11 | 348–349 | 1 | 2 | 0.080 | 4.14 | HRSCAF=293701 | bio14 (precip driest month) |
| chr11 | 359–360 | 1 | 1 | 0.072 | 4.83 | HRSCAF=219368 | bio14 (precip driest month) |
| chr11 | 441–442 | 1 | 3 | 0.087 | 5.22 | HRSCAF=219368 | bio14 (precip driest month) |

